# OmicsFM brings proteomics into the foundation model era

**DOI:** 10.64898/2026.08.25.747021

**Authors:** Sander Heyndrickx, Ralf Gabriels, Harikrishnan Ramadasan, Lennart Martens, Tine Claeys

**Author notes:** To whom correspondence should be addressed:, Address: FSVMII, Technologiepark 75, 9052 Ghent, Belgium.

## Abstract

While foundation models have been shown to learn biological representations from large transcriptomic atlases, it remained unknown whether proteomics data allow the same. We here therefore introduce OmicsFM, a modality-agnostic transformer pretrained through masked abundance reconstruction on an unprecedented proteomics data corpus of 48,837 quality-filtered proteomics profiles from 1,397 reprocessed PRIDE projects. Interestingly, despite training on 14- to 93-fold fewer profiles than matched bulk- and single-cell transcriptomic models, respectively, our proteomics model rivals both. On held-out projects, OmicsFM attention networks recovered more molecular relationships than co-expression methods and existing single-cell foundation models across nine reference databases that reveal pathway-level organization. Sample-level embeddings preserved biological structure across independent studies, and its representations transferred successfully to cell-type classification, gene-essentiality prediction, and perturbation-response prediction, while consistently outperforming task-specific models. Moreover, our results show that proteomics and transcriptomics representations capture complementary biology. OmicsFM thus firmly establishes the possibility of training highly performant proteomics-based foundation models, and their importance in modelling and uncovering fundamental biology.

## 1 Introduction

Large-scale single-cell transcriptomics atlases containing millions of human cells have enabled foundation models such as scGPT^1^, Geneformer^2^ and scPRINT^3^ to learn representations from molecular abundance data through masked gene-expression reconstruction. These representations have supported cell-type annotation, clustering, integration, gene-network inference and in-silico perturbation prediction. This approach has recently been extended to bulk transcriptomics^4^, establishing self-supervised abundance modelling as a powerful strategy for learning biological representations from large datasets.

Despite these advances, these foundation models remain exclusive to transcriptomics, only providing a single layer of cellular regulation. Indeed, between a transcript and its active protein lie translation, folding, post-translational modification, trafficking, complex assembly and degradation^5^, so transcript abundance is often an unreliable proxy for the proteins that execute cellular processes. Proteomics closes this gap by measuring protein abundance directly, offering a view of biology that is complementary to, rather than redundant with, transcriptomics. Yet no founda-tion-model framework has been established for proteomics, so whether self-supervised pretraining can learn meaningful biological representations from protein abundance data remains unknown.

Testing this requires data at scale. For transcriptomics, ARCHS4^6^ and the CELLxGENE Census^7^ already provide accessible, metadata-rich expression matrices. For proteomics, comparable large-scale data exists only in principle in the public domain, but not in practice. Three barriers have prevented such a proteomics resource from being built. First, data readiness: proteomics data are generally deposited as raw spectra requiring large-scale searching and quantification before protein abundances can be obtained. Second, data annotation: sample metadata are inconsistently structured and dispersed across repository records, filenames and publications, and typically require manual reconstruction at the run level. Third, technical heterogeneity: differences in sample preparation, fractionation, labelling, acquisition and instrumentation introduce variation that is often confounded with biology and cannot be removed indiscriminately.

In this work, we address these barriers and present OmicsFM, to our knowledge the first foundation model pretrained through self-supervised abundance reconstruction on large-scale proteomics data. We reprocessed 1,397 public PRIDE projects to assemble 48,837 quality-filtered protein abundance profiles and reconstructed their run-level metadata using an agentic extraction pipeline adapted from HAMLET^8^. The same architecture was then pretrained independently on proteomics, on bulk transcriptomics profiles from ARCHS4 and on single-cell transcriptomics profiles from CELLxGENE Census. The three models share architecture, parameter count and training configuration to enable direct comparison of the representations learned from each modality.

We evaluated the resulting representations on three levels: (i) whether feature-level attention recovers established molecular relationships and context-dependent pathway organization; (ii) whether sample-level representations preserved biological structure across projects; and (iii) whether they transfer to cell-type classification, gene-essentiality prediction and perturbation-response prediction.

Together, these results establish self-supervised abundance reconstruction as a transferable strategy beyond transcriptomics and reveal complementary biological information in proteomic and transcriptomic data.

## 2 Results

### 2.1 Three matched foundation models from proteomics, bulk and single-cell transcriptomics

To compare representations learned across the three modalities from within a matched modelling framework, we assembled a large pretraining corpus for each modality. The two transcriptomics corpora comprised 680,216 quality-filtered bulk profiles from ARCHS4^6^ and 4,550,106 cells from the CELLxGENE Census^7^, spanning 722 cell types. Because no comparable ready-to-train proteomics resource existed, we reprocessed 1,397 public PRIDE^9^ projects to assemble 48,837 quality-filtered protein abundance profiles from 1,143 projects. Each corpus was divided into training, validation and test sets at the project level (90/5/5), so no ensuring that held-out project was seen during pretraining. For the proteomics corpus, we reconstructed run-level metadata by integrating curated SDRF files, filename-pattern matching and an agentic extraction pipeline operating on repository descriptions and associated open-access publications (Methods 5.1). These annotations provide the biological labels used in the evaluations below, with coverage summarized in Table S1.

We developed OmicsFM, a modality-agnostic pre-norm transformer for heterogeneous molecular abundance profile reconstruction. OmicsFM builds on the attention masking, abundance binning and dual reconstruction objective of scGPT^1^, while incorporating sequence-informed feature representations inspired by UCE^10^ encoding feature identities using protein-language-model embeddings. Each profile is a sequence of detected features whose tokens combine a feature-identity embedding and a rank-based abundance-bin embedding. A Sample Summary Token (SST) is prepended to the sequence to aggregate sample-wide information into a fixed-dimensional representation.

During self-supervised pretraining, the model masks a subset of feature abundances and reconstructs them from the observed context using complementary feature-level and sample-level prediction heads. This objective learns both feature relationships and sample representations. Using this shared architecture, we pretrained models independently on each of the three corpora (Methods 5.2). All models use an identical 4.9M-parameter transformer backbone and training configuration, specified a priori without modality-specific tuning; they differ only in how feature identity is embedded. For every modality, we trained one variant with learned feature-identity embeddings and one with frozen ESM-C-derived sequence embeddings mapped through a small trainable projection, yielding six matched models. This design allows us to compare representations learned from the three corpora and isolate the contribution of sequence-derived feature information.

The complete workflow is summarized in Fig. 1. The analyses below examine three outputs of pretraining: feature-level attention maps, sample-level SST embeddings and learned feature-identity embeddings.

**Figure 1.**
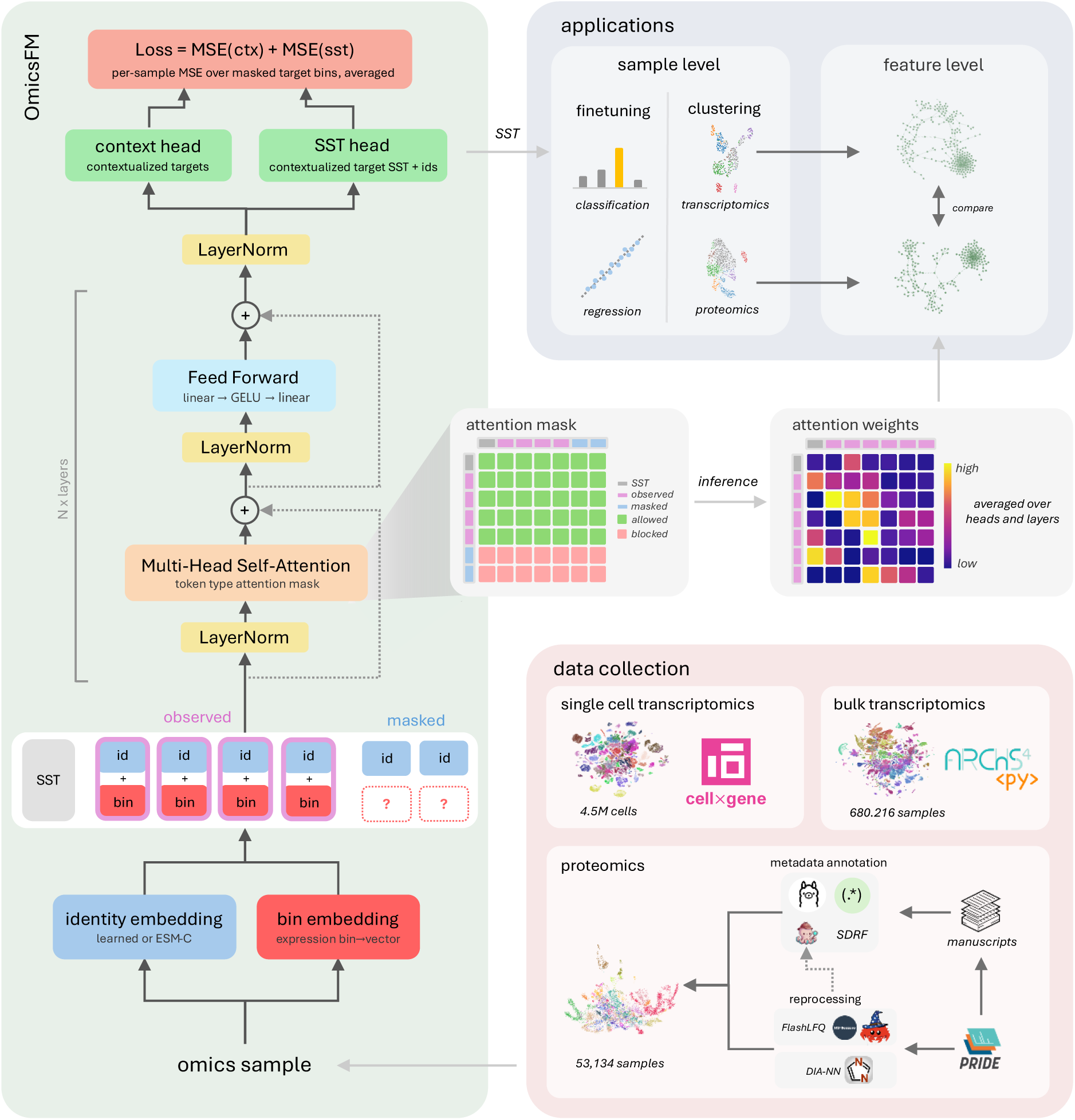
Overview of OmicsFM, a modality-agnostic foundation model for molecular expression profiles. We assemble three large training corpora (bottom right): bulk transcriptomics from ARCHS4, single-cell transcriptomics from the CELLxGENE Census, and a proteomics corpus we build by a large-scale reprocessing effort of PRIDE, whose run-level metadata we reconstruct through an automated annotation pipeline. The resulting feature-level attention, learned identity embeddings and Sample Summary Token (SST) representations were evaluated for molecular association recovery, context-dependent biological organization, sample-level structure, robustness and downstream transfer.

### 2.2 OmicsFM attention strongly recovers known molecular relationships across modalities

To reconstruct masked abundances during pretraining, OmicsFM must exploit statistical dependencies between molecular features. If these dependencies are reflected in its attention maps, functionally related features should preferentially attend to one another. Previous work has shown that attention can recover transcription factor–target relationships^1,3^. We therefore asked whether the learned attention structure recovers known molecular relationships across modalities, and additionally whether the signal extends beyond transcriptional regulation to physical interactions, protein-complex membership, pathway co-membership and shared cellular functions.

To answer this, we ranked each feature’s candidate partners by their pairwise attention scores and measured the enrichment of curated relationships among its top-ranked partners against nine reference databases: CORUM^11^, BioPlex^12^, HuRI^13^, STRING^14^, Reactome^15^, KEGG^16^, GO:CC, GO:BP^17^ and OmniPath^18^ (Methods 5.3). On held-out test projects, the attention networks derived from OmicsFM achieved the highest enrichment across all nine databases in all three modalities, an ordering that was stable across ten resampled draws of 1,000 test observations (Fig. 2A, Fig. S1). Moreover, although OmicsFM operated zero-shot, it outperformed the methods fitted directly to the benchmark data such as FAVA^19^, Pearson, GENIE3^20^, and DeepSEM^21^and substantially exceeded pre-trained single-cell foundation models, among which scGPT ranked second.

**Figure 2.**
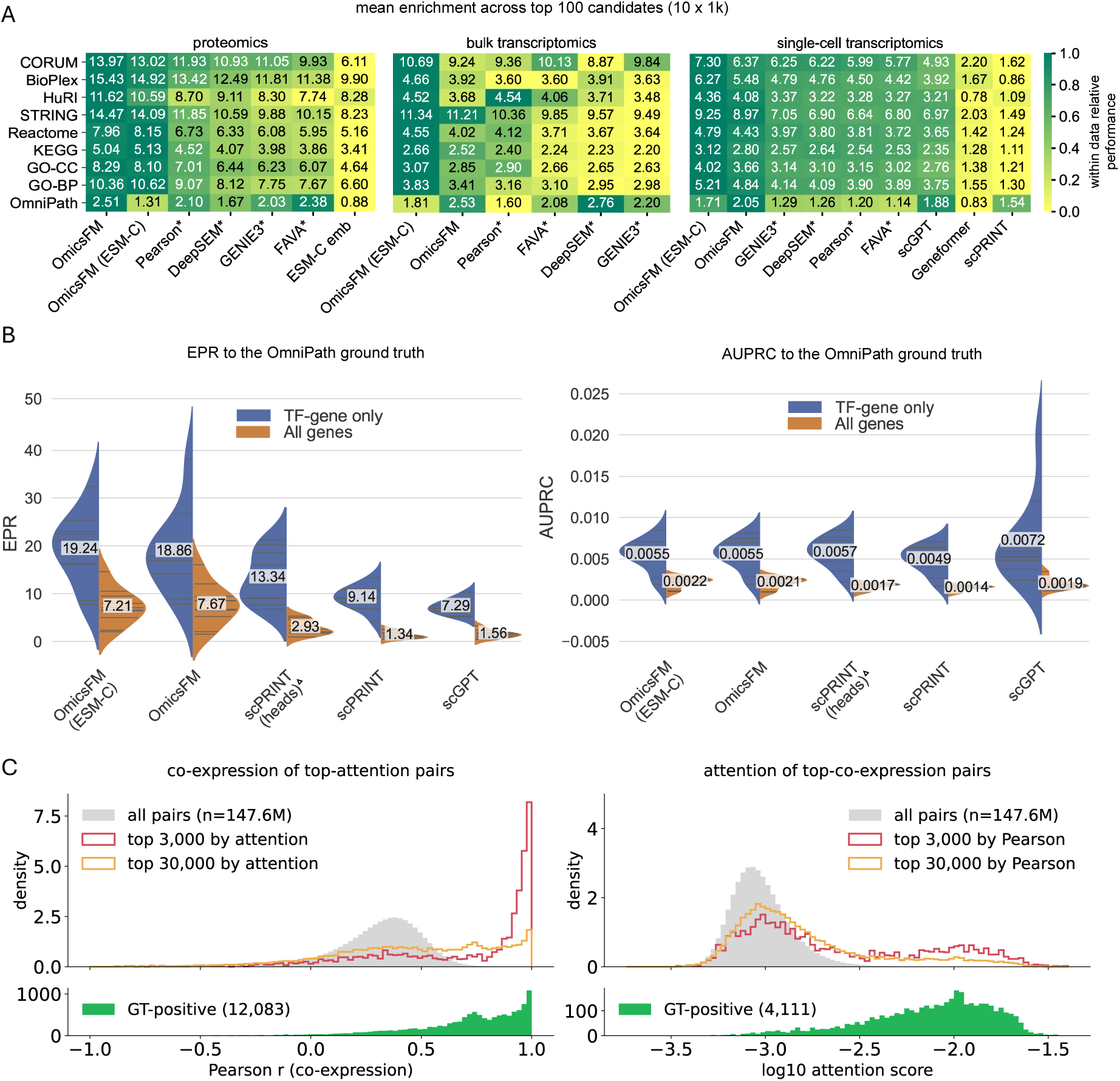
OmicsFM attention recovers biologically meaningful feature relationships across omics modalities. **(A)** Project-held-out association-network benchmark for proteomics, bulk- and single-cell transcriptomics. Ten samples of 1,000 observations were drawn with replacement from each project-disjoint test set. Within each replicate, all methods were evaluated on the same 1,000-feature universe and nine reference databases (Methods 5.3). Heatmap colors are normalized within each database and are therefore not comparable across columns. Methods marked with an asterisk* were fitted or calculated directly on each held-out replicate. **(B)** OmicsFM (single-cell transcriptomics) on the scPRINT BenGRN benchmark. Split violin plots show the Early Precision Ratio (EPR; left), which measures the enrichment of positive pairs near the top of the ranking, and the Area Under the Precision-Recall Curve (AUPRC; right), which summarizes performance across the full ranking. Results are shown for both all-gene and TF-to-gene predictions across cell types, with labels indicating the mean across cell types. scPRINT (head) is marked with (Δ) because it uses labels to select the scoring head, introducing circularity. It is therefore not a truly zero-shot method. **(C)** Comparison of the attention-derived (OmicsFM ESM-C) and co-expression-derived networks in proteomics. Left: the Pearson correlation of the top 3,000 and top 30,000 attention pairs. Right: the attention scores of the top 3,000 and top 30,000 co-expression pairs. In both, the grey distribution is all pairs, serving as the background reference, and the bottom histogram shows where the positively annotated pairs within the top 30,000 fall along the x-axis.

Protein-sequence information provided an informative prior especially for the transcriptomic models: combining feature identities with ESM-C embeddings^22^ boosted enrichment scores for every reference database except OmniPath. This gain is consistent with the sequence-only ESM-C baseline, which itself recovered appreciable enrichment. In proteomics, by contrast, the sequence prior conferred little benefit. The learned-identity variant performed marginally better across most databases, and ESM-C initialization improved enrichment only for the references capturing pathway co-membership and molecular function.

Because all references were grounded in UniProt protein space, methods operating natively on genes could in principle be disadvantaged. To control for this, we repeated the single-cell benchmark entirely in the native gene universe, using an OmicsFM model trained on the unmapped gene vocabulary, re-indexed ground truths and no mapping step, under a per-cell-type sampling design (Methods 5.3). Enrichment shifted by at most 0.09 for every method and the ranking remained unchanged (Fig. S2), so the protein grounding disadvantaged neither the transcriptomic modality nor the external models.

On the external BenGRN benchmark^3^ (Fig. 2B), OmicsFM achieved the highest Early Precision Ratio (EPR) among the evaluated foundation models, whereas scGPT scored highest across the complete ranking (AUPRC). OmicsFM therefore assigns its highest attention scores to curated OmniPath relationships more reliably than the other models.

Finally, because functionally related proteins often co-vary in abundance, we asked whether attention merely captured pairwise co-expression. We examined this relationship in both directions (Fig. 2C). Most of the highly co-expressed pairs, received attention scores similar to the background distribution. Only a minority extended into the high-attention tail, within which the annotated relationships were concentrated . Conversely, the top 3,000 attention pairs were strongly shifted toward high positive correlations, although this correspondence weakened substantially when expanded to the top 30,000 pairs. This broader set nevertheless recovered many additional annotated relationships across correlations of 0.5 to 1.0. Thus, high attention frequently coincided with co-expression, particularly at the extreme of the ranking, whereas high co-expression alone was a poor predictor of high attention.

### 2.3 Attention networks are tissue-conditioned and recover pathway organization

Having established that attention recovers known molecular relationships, we next explored whether it also captures biology specific to a tissue context. We therefore derived one attention network per tissue and modality from paired deep proteomic and transcriptomic profiles of 30 human tissues^23^ (Methods 5.4). Before probing tissue specificity, we verified that these networks retain biological signal. Across tissues, both modalities remained strongly enriched for curated relationships among the top 100 ranked partners per protein and the modality differences observed in the held-out benchmark became more pronounced at this scale (Fig. S3). Specifically, for protein complexes, for example, mean CORUM enrichment reached roughly 40-fold in the proteomic networks versus 13-fold in the transcriptomic networks, whereas OmniPath transcription-factor–target enrichment remained higher in transcriptomics (approximately 2-fold versus 1.4-fold).

With these tissue-specific networks in hand, we asked whether the model reorganizes molecular relationships according to tissue identity or instead applies a common relational structure to every input: if attention reflected a static map, its patterns would be identical across tissues. We therefore tested whether proteins within tissue-specialized pathways attend to one another more strongly in their corresponding tissue than in tissues where those pathways are inactive (Methods 5.4). Across the 30 tissues, tissue-specialized pathways showed significantly elevated within-pathway attention precisely in their expected tissues, in both proteomics (Fig. 3A, S4, S5) and transcriptomics (Fig. S4, S6): synaptic pathways in brain, secretory pathways in pancreas and salivary gland, adaptive-immune pathways in lymphoid tissues, and muscle-related cytoskeletal pathways in muscle. By contrast, broadly active housekeeping processes showed no tissue specificity, and pathway detectability across tissues largely did not account for the observed differences (Fig. S4). Attention therefore is conditioned on the biological context of the input rather than applied as a fixed relational map.

**Figure 3.**
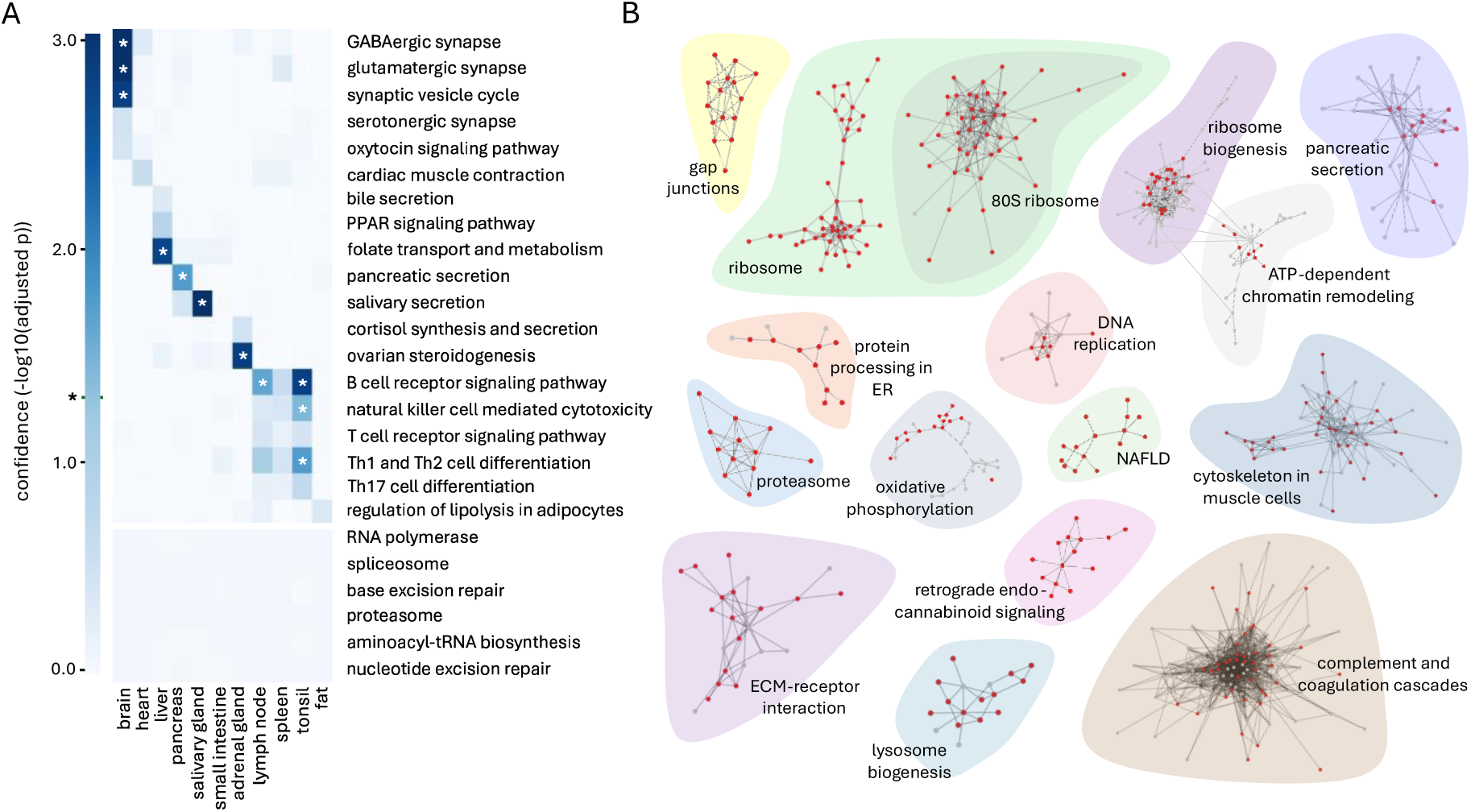
OmicsFM attention networks are tissue-conditioned and recover pathway organization across 30 human tissues. **(A)** Tissue-specificity of pathway attention across the tissues of the proteomics networks (learned identify model) (Methods 5.4), showing the significant pathway–tissue hits. Color encodes confidence as the −log10 adjusted p-value, and the marked level on the color bar with an asterisk* indicates the significance threshold (adjusted p ≤ 0.05). **(B)** Collection of protein communities detected in the proteomics attention network that are significantly enriched for KEGG pathways (Methods 5.4). For each community the full network is shown, where red nodes mark proteins belonging to the pathway assigned to that community and grey nodes mark community members outside it. Notably, some communities consist exclusively of pathway members.

**Figure 4.**
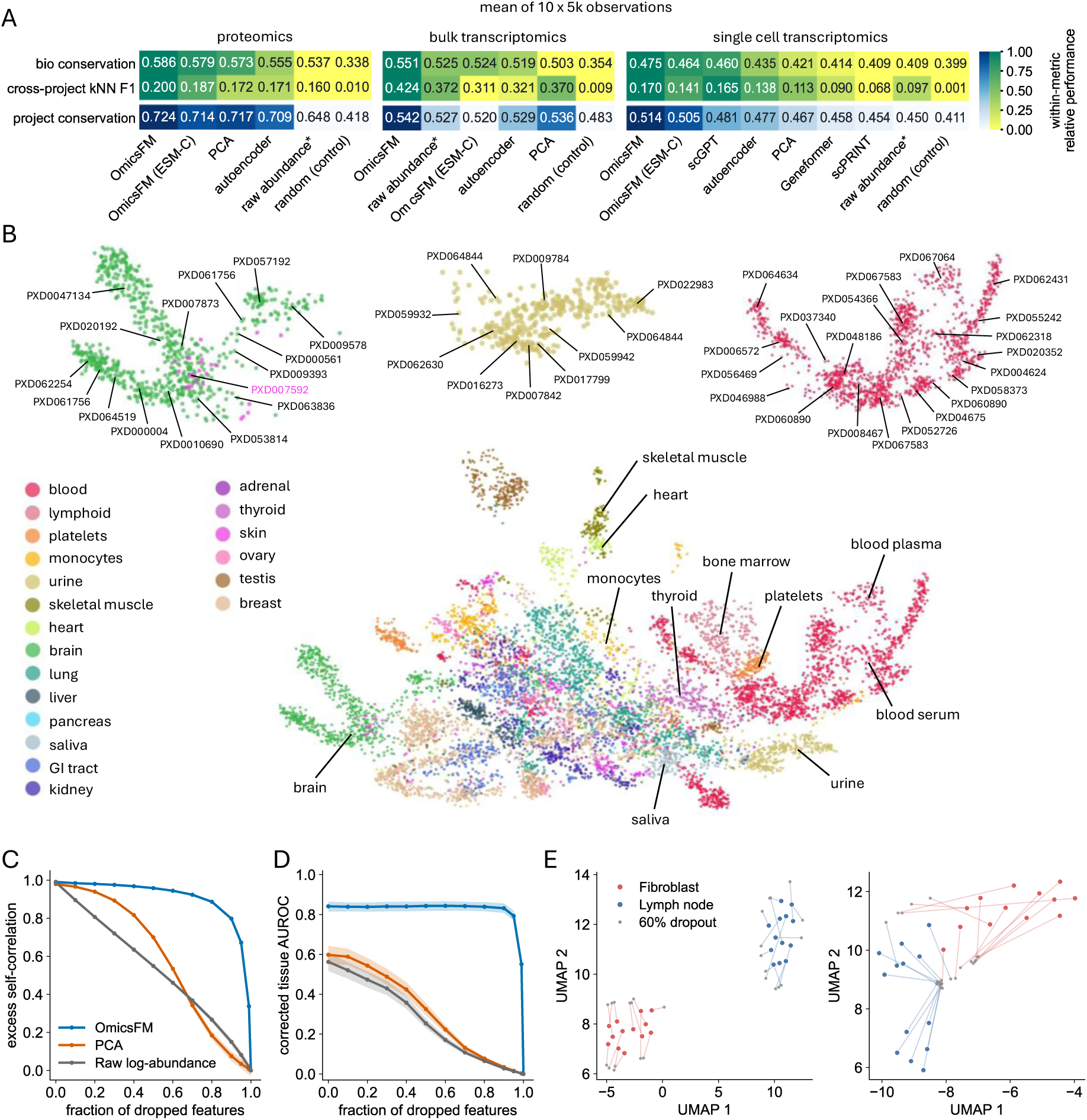
OmicsFM sample-level embeddings capture biological organization across projects and remain robust to protein dropout. **(A)** Performance of sample-level representations in proteomics, bulk transcriptomics and single-cell transcriptomics. Values are means across ten label- and project-balanced cohorts of 5,000 observations. Bio conservation summarizes six scIB^24^ metrics measuring biological organization within each corpus. Cross-project kNN F1 measures biological-label transfer when all neighbors from the query sample’s own project are excluded; because labels contributed by a single project then have no eligible neighbors and score zero in the macro-average, absolute values are deflated and scores should be read comparatively. Project conservation summarizes the degree to which project identity remains encoded in the representation, with higher values indicating stronger conservation. All learned and reduced representations were matched to the same dimensionality. Raw abundance, marked with an asterisk, is not a learned representation and retains the full approximately 20,000-feature protein space. Colors indicate relative performance within each modality and metric. **(B)** UMAP of the proteomics OmicsFM SST embeddings, colored by tissue annotation. Selected brain, urine and blood regions are enlarged above, with arrows identifying the contributing ProteomeXchange projects. Samples from several independent projects occupy the same tissue-associated regions. **(C)** Excess self-correlation between perturbed (dropout) and unperturbed representations as increasing fractions of detected proteins were removed from the test set. **(D)** Chance-corrected cross-project tissue-recovery AUROC, using unperturbed samples as references and excluding references from the query’s project (n 622 across seven tissues from the test set). In C and D, OmicsFM SST is compared with dimension-matched PCA and full-dimensional log-transformed abundance profiles; lines show means across five dropout masks and shading shows 95% bootstrap confidence intervals. **(E)** Paired UMAP projections of 13 fibroblast and 13 lymph-node samples from the test set before and after 60% dropout. Large colored points show unperturbed samples, small grey points their perturbed counterparts, and lines connect paired representations. UMAPs were fitted separately and are not directly comparable between methods.

**Figure 5.**
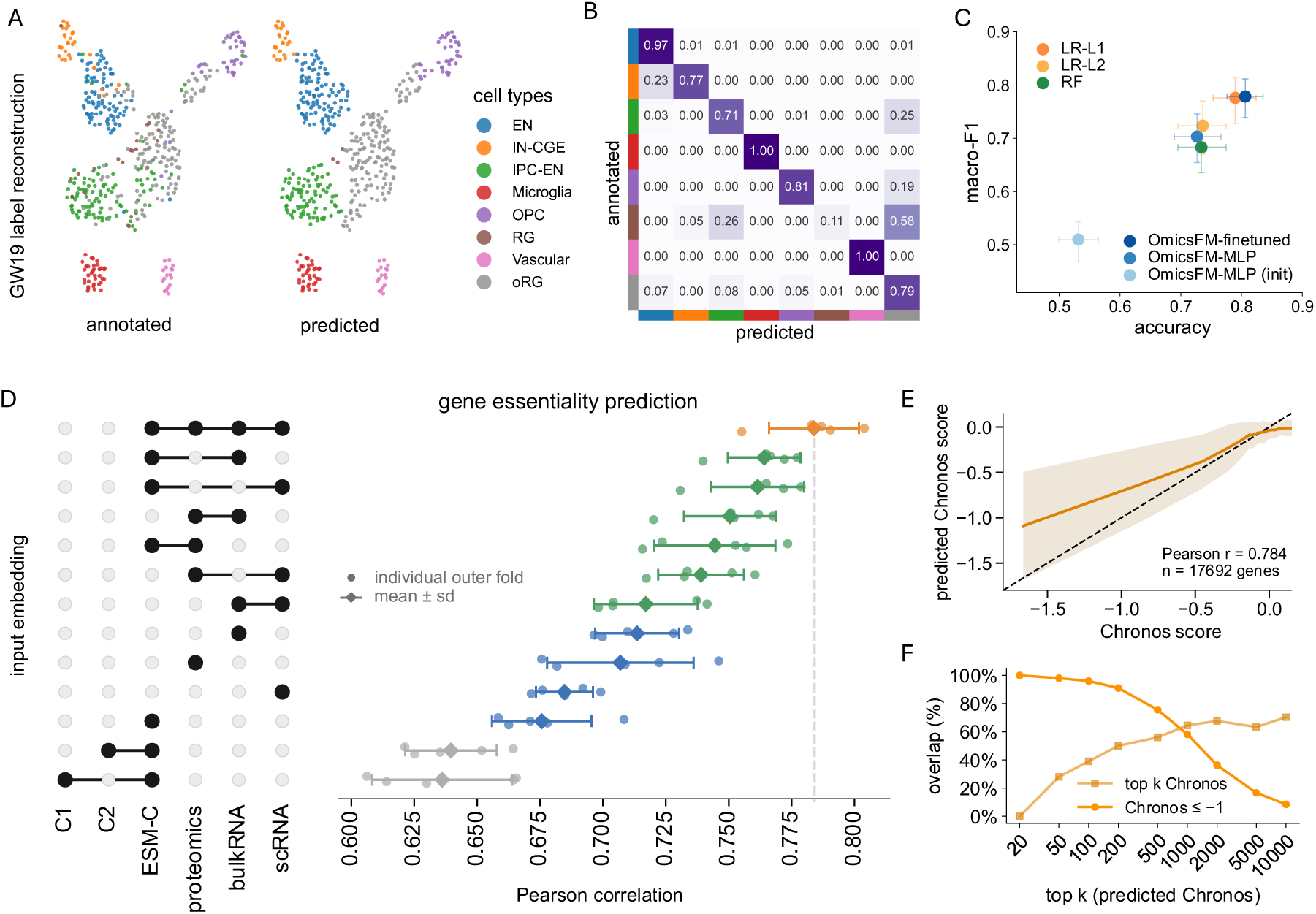
Pretrained representations transfer to diverse downstream tasks. (A-C) Cell-type classification on sc proteomics profiles. **(A)** GW19 cells in the same UMAP projection, colored by annotated and OmicsFM-predicted cell type. **(B)** Row-normalized confusion matrix for the end-to-end fine-tuned model on GW19. **(C)** Accuracy and macro-F1 on GW19 for end-to-end OmicsFM fine-tuning, MLP classifiers trained on frozen pretrained or non-pretrained (init) OmicsFM representations, and LR-L1, LR-L2, and RF classifiers trained directly on log₂(x 1)-transformed protein abundances. Points represent performance averaged across three independently seeded runs for the OmicsFM methods and single fitted models for the classical baselines. Error bars show percentile 95% confidence intervals from 1,000 bootstrap resamples of held-out GW19 cells. **(D–F)** Prediction of general gene essentiality from gene representations. **(D)** Pearson correlation between predicted and mean DepMap Chronos scores for MLPs trained on individual or concatenated frozen representations. The matrix indicates the representations supplied to each model. C1 denotes ESM-C combined with Gaussian control features, and C2 denotes ESM-C combined with gene-permuted OmicsFM embeddings. **(E)** Concordance between true and out-of-fold predicted Chronos scores for the model combining representations (orange). Genes were divided into equally sized bins according to their annotated Chronos score; the orange line and shaded region show the mean ± sd of the predictions within each bin, and the dashed line indicates identity. **(F)** Recovery of essential genes among the top-k predictions from the combined model. Curves show the percentage of predicted genes also present among the annotated top-k most essential genes and the percentage with an annotated mean Chronos score ≤ −1.

In the previous analysis, prior knowledge defined the pathways, and attention was measured within them. We next reversed the question, letting the attention network itself define groups and asking whether these correspond to known biology. Community detection on the top 3,000 attention edges, performed without any biological annotation, identified numerous protein modules significantly enriched for KEGG pathways (Fig. 3B). Several modules were highly specific, consisting almost exclusively of ribosomal proteins, gap-junction components or proteasome subunits, whereas others extended beyond their most enriched pathway yet remained biologically coherent, for example linking ribosome biogenesis to ATP-dependent chromatin remodeling. The model therefore acquires a degree of pathway-level organization during self-supervised pretraining.

### 2.4 Sample-level representations capture biological structure across modalities

Complementary to the feature-level representations, the SST is trained to compress the global abundance structure of a sample into a single fixed-dimensional embedding. Because a useful sample-level representation should group samples by their biology, we asked how much of the biological structure across samples this embedding retains in each modality, and we quantified this through three complementary properties: (i) biological conservation, the extent to which samples cluster by tissue or cell-type label; (ii) cross-project label transfer, whether a sample’s label can be recovered from its neighbors in other projects; and (iii) project conservation, the degree to which study-specific structure persists in the embedding. Each property was scored on ten label- and project-balanced evaluation cohorts per modality (Methods 5.5). As reference points, we included representations of the same data without pretraining (random, raw abundance, PCA, autoencoder), and for single-cell transcriptomics we additionally compared against scGPT, Geneformer^2^ and scPRINT^3^.

OmicsFM obtained the highest mean bio-conservation score in all three modalities (Fig. 4A, S7 and S8), ranking first in each of the ten evaluation cohorts in proteomics and single-cell transcriptomics. In bulk transcriptomics the lead was less stable, holding in eight of ten cohorts. Initializing feature identities with ESM-C did not improve sample-level representations in any modality. All external single-cell foundation models scored above both the random baseline and raw abundances, but only scGPT approached the OmicsFM representations, whereas Geneformer and scPRINT fell below PCA and the autoencoder.

OmicsFM also achieved the highest cross-project kNN F1 in each modality, so its local neighborhoods remained informative of biological identity even when all neighbors from a sample’s own project were excluded. At the same time, it obtained the highest project-conservation score, which on its own would indicate a representation dominated by study-specific variation. However, such a representation could not transfer labels across project boundaries, and the two results indicate embeddings that remain biologically aligned across studies while still encoding study-specific structure. Repeating the evaluation on the project-held-out test sets alone confirmed OmicsFM as the strongest representation in proteomics and bulk transcriptomics, with scGPT ahead in single-cell transcriptomics (Fig. S9). Here, every observation was unseen by OmicsFM, but no equivalent guarantee holds for scGPT’s larger pretraining corpus.

This biological organization is directly visible in the OmicsFM proteomics SST embeddings (Fig. 4B). Samples annotated as brain, urine and blood formed regions populated by multiple independent projects, and the broader arrangement followed known physiology: plasma and serum lay adjacent to platelets and bone marrow, and heart lay beside skeletal muscle. One group of skin-annotated samples appeared within the brain region, but these samples originated from PXD007592, a study of cerebral metastases in melanoma patients, therefore their position is biologically plausible.

Finally, because the SST is pretrained on randomly subsampled protein sets (1,024 features), we used the proteomics model and its test samples to test whether its representations remain stable when detected proteins are removed (Methods 5.5). Excess self-correlation stayed above 0.9 through 80% dropout and declined sharply only beyond 90%, whereas PCA and log-transformed abundance profiles deteriorated steadily and approached zero by 90% (Fig. 4C). Importantly, this stability preserved biological organization: cross-project tissue retrieval remained nearly unchanged through 80% dropout and stayed substantially above chance even after 95% of detected proteins were removed (Fig. 4D). The SST thus retains both sample identity and tissue-level organization from severely incomplete proteomic profiles.

### 2.5 Pretrained representations transfer to downstream tasks

To determine whether the representations learned during self-supervised pretraining transfer to new applications, we adapted the pretrained models to supervised downstream tasks, comparing end-to-end fine-tuning with predictors trained on frozen pretrained representations and with matched randomly initialized controls. We evaluated three applications: (i) cell-type classification from single-cell proteomics profiles (data from Wu ^25^); (ii) gene-essentiality prediction from the learned feature-identity embeddings (data from Tsherniak ^26^ and Dempster ^27^) and (iii) perturbation predictions (data from Replogle K562^28^, Adamson^29^ and Norman^30^) (Methods 5.6).

#### For cell-type classification

we used data from PXD071075 which was purposefully excluded from pretraining. This dataset profiles single-cell proteomes from three developing human brain donors at gestational weeks (GW) 13, 15 and 19. Models were trained on the GW13 and GW15 donors and evaluated on held-out GW19, representing a combined donor and developmental-stage shift (Fig. 5A-C). End-to-end fine-tuning reached a macro-F1 of 0.778 and an accuracy of 0.806, an MLP on frozen pretrained representations reached 0.703 and 0.727, whereas the same predictor on a frozen randomly initialized encoder reached only 0.510 and 0.531. The pretrained SST therefore carried substantial cell-type information before any task-specific optimization, and most residual errors occurred among closely related progenitor populations (radial glia, outer radial glia and intermediate progenitor cells). Nevertheless, L1-regularized logistic regression on protein abundances matched the fine-tuned model’s performance, so this experiment demonstrates that the pretrained representation transfers and improves markedly over an untrained encoder, but not that OmicsFM outperforms conventional classifiers on this marker-driven task.

#### Gene essentiality prediction

Our second application probes a different output of pretraining, namely the feature-identity embeddings, which we used to predict gene essentiality (Methods 5.6). Essentiality reflects how strongly the loss of a gene compromises cellular fitness and thus depends on the molecular functions and interactions of its product. As ground truth, we averaged each gene’s Chronos gene-effect scores from the DepMap Public 24Q4 CRISPR knockout screens across 1,178 cancer cell lines, yielding one average essentiality score for each of 17,692 genes. We then tested how well the frozen feature-identity embeddings predict these scores (Fig. 5D-F).

The proteomics and bulk-transcriptomic identity embeddings outperformed the sequence-based ESM-C representation (r 0.706 and 0.713 versus 0.674), with gains exceeding the fold-to-fold variation, whereas the single-cell embedding was only numerically higher (r 0.683). Combining modalities improved prediction further, most notably proteomics with bulk transcriptomics (r 0.751), whereas combining the two transcriptomic representations added little over either alone. Integrating all three OmicsFM modalities with ESM-C achieved the best performance (r 0.784), improving by 0.109 over ESM-C alone and by 0.035 over the best OmicsFM-only combination, a gain exceeding the fold-to-fold variation of either model. Proteomic, transcriptomic and sequence representations therefore encode complementary information that can be integrated for downstream prediction.

#### In silico-perturbation prediction

Our final application asks whether pretrained representations improve perturbation-response prediction. We evaluated on three CRISPR screens, Replogle K562^28^, Adamson^29^ and Norman^30^, comparing the three fine-tuned OmicsFM models against an identical model trained from scratch, scGPT, GEARS^31^ and a linear baseline (Fig. 6).

**Figure 6.**
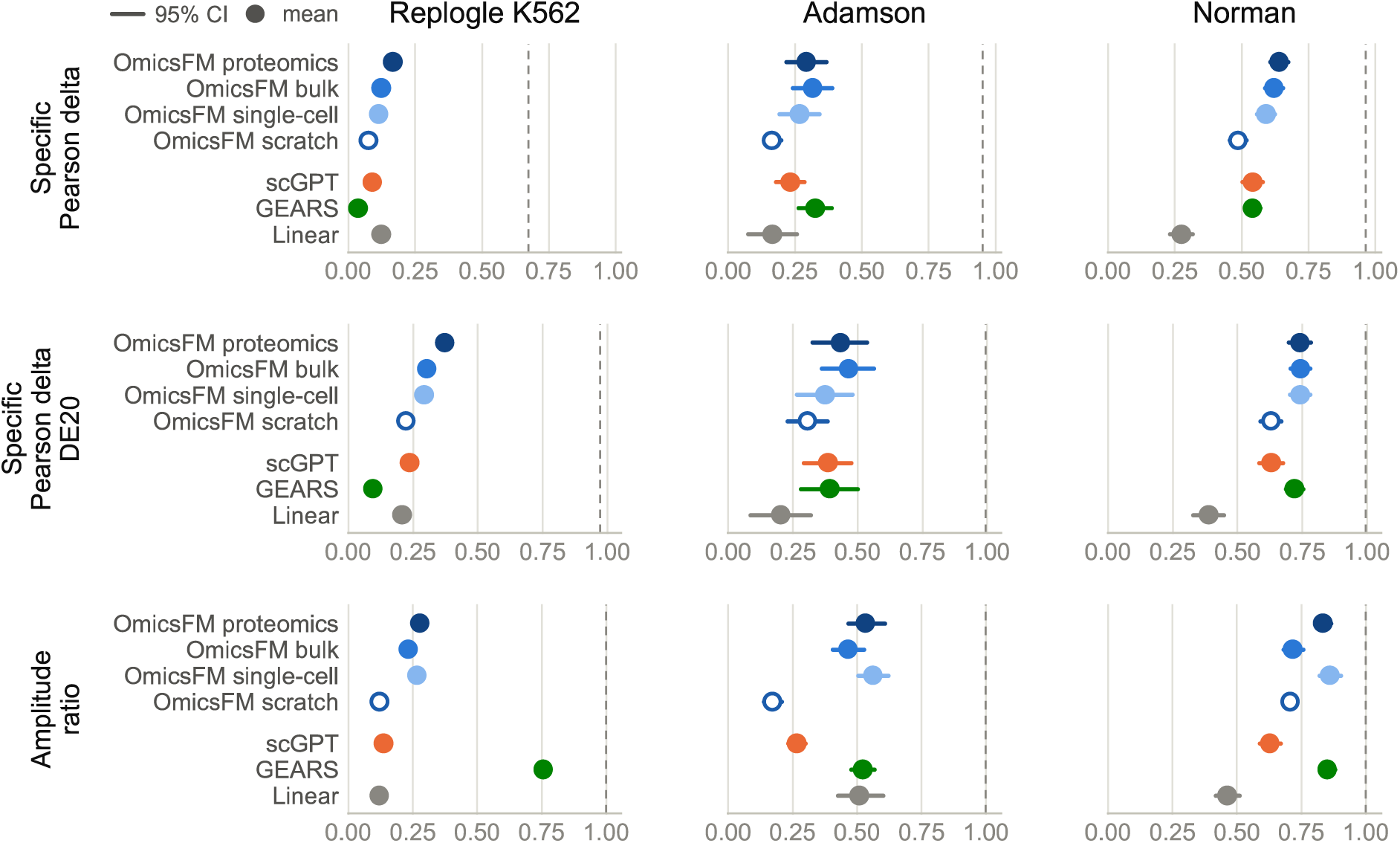
Pretrained OmicsFM representations improve prediction of perturbation-specific responses. Fine-tuned models were evaluated on three held-out CRISPR screens (columns: Replogle K562^28^, Adamson^29^ and Norman^30^) using metrics that isolate the perturbation-specific response after removing the generic response (Methods 5.7): the specific Pearson delta across all genes, the same correlation restricted to each perturbation’s twenty strongest differentially expressed genes, and the amplitude ratio (predicted over observed response magnitude; 1 fully recovered). Points show means, bars 95% confidence intervals across folds. Dashed lines give the reliability ceiling, the maximum correlation attainable against the noise of the experimental data (Methods 5.7), or exact amplitude recovery in the bottom row.

Perturbation-specific effects proved partially predictable. Sample wide Specific correlations were modest on Replogle and Adamson but rose substantially when restricted to the twenty strongest differentially expressed genes (DE20), and were markedly higher on Norman throughout. On DE20, every OmicsFM model and scGPT exceeded the linear baseline on all three datasets. Crucially, the pretrained OmicsFM models outperformed their from-scratch counterpart across all metrics and datasets, directly demonstrating that the pretrained representations carry information relevant to predicting perturbation effects. The OmicsFM models also consistently matched or exceeded scGPT, in line with their stronger recovery of known gene–gene relationships, and the proteomics model was again among the strongest across the board despite being modality misaligned and its pretraining corpus being orders of magnitude smaller and of a different modality than the fine-tuning data. GEARS behaved inconsistently, competitive on Adamson and Norman but near the bottom on Replogle. All models, however, systematically underestimated the magnitude of perturbation-specific responses, with amplitude ratios below one throughout.

## 3 Discussion

### 3.1 The foundation-model paradigm transfers beyond single-cell transcriptomics

In this work, we establish that the foundation-model paradigm transfers to proteomics. OmicsFM is, to our knowledge, the first such model and its representations carried biological information at every level we examined, from feature-level attention to sample and gene-level embedding. This is notable given the proteomics corpus’ heterogeneity and smaller size relative to the transcriptomics corpora. Although the representations are not exclusively biological, since the sample embeddings also retained study-associated structure, they consistently captured biology that transferred across datasets and tasks. The matched transcriptomics models confirmed that the same modelling principle holds generally by recovering the structure expected of single-cell foundation models and extending it to tissue-level RNA profiles, consistent with the recently introduced BulkFormer^4^. These transcriptomics OmicsFM models served as positive controls for the architecture and as matched reference points for interpreting the proteomics OmicsFM model. Together, the results establish self-supervised abundance reconstruction as a general representation-learning strategy for molecular profiles, thus bringing proteomics firmly into the foundation model era.

### 3.2 Proteomics models rival their transcriptomic counterparts from far less data

Within this reference framework, the most striking result was how little data the proteomics model required to reach performance comparable to its transcriptomic counterparts. Indeed, pretrained on roughly 14-fold and 93-fold less data for bulk and single-cell transcriptomics respectively, it rivalled both throughout the study across attention-based molecular relationships, sample-level biological conservation and fine-tuned perturbation results. Despite the modality mismatch between pretraining and fine-tuning, the proteomics model transferred successfully to RNA perturbation prediction, suggesting that it captured reusable biological relationships rather than mass-spectrometry-specific patterns. However, comparable performance did not imply interchangeable representations. Proteomics and transcriptomics attention recovered different classes of interactions. The same complementarity emerged in gene-essentiality prediction where combining proteomic and transcriptomic feature-identity embeddings outperformed the individual modalities. This was further improved by adding the sequence-derived ESM-C^22^ representation. By contrast, combining the bulk- and single-cell-transcriptomic embeddings added little over bulk transcriptomics alone. These comparisons cannot establish that proteomics is intrinsically more data-efficient, because modality and corpus size remain confounded. They do show, however, that far less data sufficed to match considerably larger transcriptomic corpora, which raises the broader question of how strongly representation quality depends on model size and the scale of the pretraining corpus.

### 3.3 The benefits of scaling model parameters and pretraining corpora remain unclear

The external single-cell foundation models provided an additional calibration for the impact of scale. Because these external models differ from OmicsFM in architecture, objective and training corpus, we used them only to confirm that our transcriptomic models were competitive. Unexpectedly, our single-cell model generally matched or exceeded these substantially larger models despite having fewer parameters and being pretrained on far less cells. Although this comparison remains confounded by architectural differences, our results do not support the notion that increasing model or dataset size alone yields more biologically informative representations.

The wider literature on scaling laws remains similarly unresolved. Broad evaluations by DenAdel found little benefit from expanding pretraining corpora^32^, whereas scaling behavior has been reported for Geneformer^2^ on fine-tuned gene-level tasks^33^. However, this negative evidence is weakened by two problems. First, experimental design: DenAdel grow the pretraining corpus while model capacity stays fixed, whereas true scaling laws require data, parameters and compute to grow together^34^; a data-only sweep can therefore miss real scaling. Second, the endpoints themselves fail as measuring instruments in two opposite ways. Cell-type classification suffers from a ceiling: annotation is essentially marker recognition, solvable from simple linear relationships between a few marker genes. Boiarsky ^35^ accordingly showed that logistic regression matches fine-tuned foundation models even in few-shot settings, and it likewise matched our end-to-end fine-tuned model on single-cell proteomics. Because these labels are marker-derived, the benchmark likely cannot reward biological structure beyond what marker logic encodes. Conversely, perturbation prediction suffers from a hidden floor: its metrics are dominated by variance that all conditions share, namely baseline expression in absolute metrics and the generic perturbation response in delta metrics^36,37^, so that predicting the mean training-set response for every perturbation already matches fine-tuned foundation models^38^. These benchmarks therefore measure the generic response rather than the response specific to the perturbation under test, which is the biologically interesting quantity. Our own perturbation experiments address this by removing the generic response and only scoring the perturbation-specific component. Here, every pretrained OmicsFM model outperformed both the linear baseline and an identically configured model trained from scratch, across all three screens and all metrics. Although this performance remained far from perfect, it shows that the apparent parity between foundation models and simple baselines on this task is a property of the metric rather than of the models. In both cases, the score moves for reasons unrelated to representation quality. Resolving the question of scaling law for omics foundation models will require in-depth experimentation. This is essential as the answer will determine whether further progress primarily requires greater scale, or rather improvements in data quality, objectives and architecture.

### 3.4 The current practical utility of omics foundation models

Regardless of how the scaling question resolves, today’s models already provide capabilities that conventional methods lack. Attention networks can act as hypothesis engines for protein relationships, covering the whole proteome and carrying signal beyond co-expression and tissue context. Indeed, the strongest partners of a poorly characterized protein offer candidate functions by association, and a strongly filtered attention network can serve as a data-derived scaffold for network-based analyses. The sample embeddings are stable and biologically informative to support neighbor search, out-of-distribution screening, and comparing profiles with very different numbers of detected proteins. The feature-identity embeddings, finally, complement sequence-derived embeddings such as ESM-C and can serve as a building block for other models, much as ESM-C served us here. However, these representations are not universally preferable to task-specific models. When a supervised endpoint can be resolved from a small set of discriminative markers, pretraining and fine-tuning may offer little advantage: in our cell-type classification experiment, logistic regression matched the fine-tuned model, making the simpler method the more appropriate choice, and similar considerations are likely to apply to other marker-driven tasks. The distinctive value of pretrained representations instead emerges when the relationships among features are themselves part of the object of study.

### 3.5 Limitations and outlook

The practical value of these representations is nevertheless bounded by several constraints. Although the modelling framework was matched, the three corpora differed in size, composition, feature coverage and experimental heterogeneity, so the comparison reflects what current public data allow rather than an intrinsic ranking of modalities. We also retained project effects, as technical variation is strongly confounded with biology across studies; the sample embeddings should consequently not be interpreted as batch-integrated representations, despite scoring strongest on biological transfer across projects among the tested representations.

The current proteomics corpus captures only part of the information accessible to the modality. Searches excluded isoforms or post-translational modifications, the data were predominantly bulk and shallow, and protein groups were excluded to maintain a comparable feature space across modalities. Additionally, the run-level metadata were reconstructed rather than curated; although agentic annotation enabled corpus assembly at this scale, its incomplete coverage and accuracy may introduce additional uncertainty into label-based analyses. Addressing this will require rich, standardized proteomics atlases with greater cellular resolution, systematic perturbations and proteoform-level measurements accompanied by standardized metadata and calibration samples. Indeed, the reprocessing and metadata reconstruction required here show how much reusable information remains inaccessible in current public repositories.

Beyond the corpus, the evaluations themselves constrain what can be concluded. The attention-network benchmark measures whether a model encodes molecular relationships but also how readily those relationships can be extracted. Furthermore, the reference databases are likely incomplete and biased toward well-studied proteins, rewarding known biology without validating novel edges. Because attention was symmetrized to match these references, its potential directionality remains untested, and no physical or causal interpretation is implied.

Looking ahead, modality complementarity, currently exploited only by concatenating representations after pretraining, could instead be captured through joint multimodal training. This would allow proteomic and transcriptomic information to shape one another while retaining modality-specific input handling, including protein groups and, ultimately, isoforms and post-translational modifications. Whether such alignment requires paired measurements or can be learned from disjoint corpora connected through shared gene and protein identities will determine how much paired multi-omics data needs to be generated.

If these gaps close, the complementarity already visible between modalities may deepen further, surfacing regulatory relationships that neither proteomics nor transcriptomics can reveal alone.

## 4 Methods

### 4.1 Data collection and annotation

#### Proteomics

**spectral data reprocessing.** Public proteomics datasets were retrieved from the PRIDE Archive^9^ and reprocessed from the deposited raw files. Each project was routed by its acquisition mode (DDA or DIA) to one of two pipelines, in which spectra were searched against the UniProt^39^ human reference proteome (canonical sequences, no isoforms; release 2022-03-31), supplemented with the cRAP^40^ set of common contaminants.

For DDA, raw files from 1,199 projects were converted to mzML and searched with Sage^41^ (v0.14.7) against an internally generated target-decoy database (reversed sequences). Tryptic digestion (no cleavage C-terminal to proline) was specified with one missed cleavage, peptide length 5–50 residues, peptide mass 500–5,000 Da and precursor charges 2-4; oxidation of methionine and carbamidomethylation of cysteine were specified as variable modifications for every project. PSMs were rescored with MS²Rescore^42^ (v3.2.0), and identifications were retained at 1% run-level FDR. Proteins were then assembled into groups by parsimony, merging proteins indistinguishable by their observed peptides, and protein-group abundances were quantified label-free with FlashLFQ^43^ (v1.2.6; 10 ppm peak tolerance, ≥2 isotope peaks, match-between-runs disabled).

For DIA, runs from 230 projects were analyzed with DIA-NN^44^ (v2.2.0) in library-free mode. Digestion settings were identical for library prediction and search: tryptic cleavage, one missed cleavage, peptide length 7–30, precursor m/z 300–1,800, fragment m/z 200–1,800, precursor charges 1–4 and N-terminal methionine excision. Carbamidomethylation of cysteine was fixed and oxidation of methionine variable (≤1 per peptide). Precursors were filtered at 1% q-value, proteins were grouped at the protein-name level with group-level FDR controlled at 1% q-value, and abundances were taken from DIA-NN’s protein-group output.

In total across DDA and DIA, 78,601 spectral data files were searched from 1,397 projects.

#### Proteomics: metadata annotation

Each run was annotated with 17 fields spanning sample biology (organism, tissue, disease, cell part, cell line), sample preparation (enzymes, modifications, labelling, fractionation, enrichment) and MS configuration (instrument, fragmentation, collision energy, acquisition mode, LC column, gradient time, ionization). Values were drawn from three sources and merged with priority-based conflict resolution and per-field evidence tracking.

First, experimental details were extracted from associated publications using an adapted HAMLET^8^ pipeline. For each project, linked open-access full texts and dataset descriptors were retrieved through the PRIDE Archive REST API, the NCBI ID Converter and the BioC API (with Europe PMC abstracts as fallback) and passed to a locally served large language model (LLM) (Qwen3.5-35B-A3B, 4-bit quantized, via Ollama; 32k context) with a fixed instruction set to annotate the desired metadata fields in a structured JSON schema per project. Second, where available, Sample and Data Relationship Format (SDRF) files were downloaded from the PRIDE Archive REST API and their column headers mapped to our metadata schema. Because these SDRF files are community curated, they were treated as the highest-confidence source. Third, a library of 300 regular-expression patterns extracted instruments, fragmentation methods, organisms, tissues, cell lines, acquisition modes and labelling strategies directly from run filenames. A first-match-wins rule was applied except for inherently multi-valued fields such as search modifications, with domain-specific priorities where appropriate (for example, when a run name referenced both a biofluid and a solid organ, the biofluid was preferred as the more likely analyzed sample). Additionally, MLMarker^45^ was used to predict tissue type from the abundance profiles as a sanity check.

Free-text values from the LLM and SDRF sources were standardized against controlled vocabularies using a SapBERT encoder^46^ (cambridgeltl/SapBERT-from-PubMedBERT-fulltext) with nearest-neighbour search: each value was mapped to its closest vocabulary term and retained when their cosine similarity exceeded 0.7. Field-specific vocabularies were drawn from NCBITaxon^47^ (organism), UBERON^48^ (tissue), the Cell Ontology^49^ (cell type) and MONDO^50^ (disease), and a small set of post-normalization rules collapsed over-specific terms to the intended granularity (for example, “triple negative breast cancer” → “breast cancer”). For each field, values from the three sources were then merged under a fixed priority order (SDRF > filename > LLM), selecting the highest-priority non-empty value and pairing it with an evidence annotation recording which sources contributed and whether they agreed, yielding a single run-level metadata table with transparent provenance.

The manuscript-extraction pipeline was validated against manually curated ground-truth annotations for 30 manuscripts used during development. To confirm generalization beyond this set, extractions for a further 30 manuscripts, excluded from development, were evaluated in the same way, with extracted fields scored by precision, recall and F1 (Table S2).

#### Proteomics: filtering and splitting

Starting from 78,601 reprocessed runs spanning 1,397 ProteomeXchange projects, the corpus was assembled through successive filtering steps. First, because 32 projects contained both DDA and DIA runs and were therefore processed by both pipelines, 3,225 runs appeared in both profiles. Each was resolved to the profile matching its annotated acquisition mode where that mode was unambiguous, while runs annotated as both DDA and DIA were discarded, dropping 5,331 rows and leaving 73,270 runs from 1,390 projects. Second, fractionated runs were identified from the fraction annotation and aggregated by summing protein quantities across fractions, collapsing 6,916 fraction runs into 708 samples and yielding 67,062 samples. Third, a detection filter requiring at least 500 quantified proteins removed 18,225 low-coverage samples (27.2%), retaining 48,837 samples from 1,143 projects. Finally, the corpus was split into training, validation and test sets in a 90/5/5 ratio by project count (ProteomeXchange identifier), so that all samples from a study remain within a single split, yielding 45,188 training samples (1,028 projects), 1,568 validation samples (57 projects) and 2,081 test samples (58 projects).

#### Bulk transcriptomics

Bulk RNA-sequencing profiles were obtained from ARCHS4^6^. Single-cell experiments were excluded using ARCHS4’s per-sample single-cell probability (threshold 0.5), retaining 773,799 of 1,093,742 candidate samples. Quality control requiring at least 5,000 detected genes, at least 1,000,000 total mapped counts and non-empty source metadata left 680,216 samples across 67,186 gene symbols. To place this modality in the same feature space as the proteomics models, gene-level counts were projected into UniProt^39^ protein space by summing counts over all gene symbols mapping to a given accession. 21,191 gene symbols (31.5%) mapped, covering 18,608 of the 20,272 reference proteins. The corpus was split into training, validation and test sets in a 90/5/5 ratio at the project level (GEO series), yielding 614,169 training, 34,945 validation and 31,102 test samples.

#### Single-cell transcriptomics

Single-cell RNA-sequencing profiles were obtained from the CELLxGENE Census^7^. To ensure broad and balanced coverage of cellular identities, sampling was stratified across 27 anatomical tissues, with up to 200,000 cells drawn per tissue; within each tissue, cells were sampled with probability inversely proportional to cell-type frequency, oversampling rare populations. Only primary, cell-suspension data were retained, and profiles passed conventional single-cell quality control, requiring at least 200 detected genes and 500 total UMIs per cell and a mitochondrial read fraction below 20%. The resulting corpus comprised 4,550,106 cells spanning 722 annotated cell types, drawn from 298 datasets and 6,905 donors, with each cell retaining its original CELLxGENE tissue and cell-type annotations. The corpus was split into training, validation and test sets in a 90/5/5 ratio at the project level (CELLxGENE dataset), yielding 3,546,382 training, 361,037 validation and 642,687 test cells.

### 4.2 OmicsFM architecture and pretraining

#### Architecture overview

OmicsFM is a self-supervised pre-norm transformer that learns representations of molecular expression profiles by reconstructing masked abundance measurements (Fig. 1): within each sample, a subset of abundances is withheld and predicted from the remaining observed features, which requires the model to learn how features co-vary within biological samples. Its design consolidates components established in single-cell foundation models, adopting the rank-based abundance binning, token-type attention masking and dual reconstruction objective of scGPT^1^ and the protein-language-model feature identities of UCE^10^, within a deliberately modality-agnostic architecture that allows matched models to be pretrained on each modality and compared directly.

#### Tokenization

Within each sample, detected features are ranked by abundance and partitioned into B equal-frequency bins, ordered from bin 1 (lowest) to bin B (highest), while undetected features receive bin 0. This rank-based binning is robust to the wide dynamic range of expression data and to technical variation across projects^1^, a primary consideration for corpora assembled from thousands of independent studies. For proteomics, protein groups containing multiple indistinguishable proteins were excluded from the model input, so that each abundance is assigned to an individual protein. Each feature is represented by a single token, embedded as the sum of a feature-identity embedding and an abundance-bin embedding. The identity embedding is either learned or initialized from frozen ESM-C^22^ embeddings projected to the model dimension through a trainable multilayer perceptron, and these two options define the two variants trained per modality.

#### Input construction

For each sample, an input sequence is assembled by subsampling up to a fixed number of detected features, partitioned into observed and masked tokens. Both carry the feature-identity embedding. Observed tokens add the abundance-bin embedding, whereas masked tokens omit it, and these withheld bins are the reconstruction targets. A Sample Summary Token (SST), analogous to the commonly used CLS token, is prepended so that its contextualized representation summarizes the sample. Attention is restricted by a token-type mask (Fig. 1): only the SST and the observed tokens act as attention keys, so every token attends to the SST and the observed set, while masked tokens are visible to no position, not even themselves. Information therefore flows from observed abundances into masked positions and never back. Consequently, no masked position sees another, making predictions invariant to which features are masked alongside them.

#### Learning objective

Masked abundance reconstruction couples two prediction heads: the feature-level head predicts each masked abundance bin from the contextualized embedding of the corresponding masked token, and the sample-level head predicts the same bin from the contextualized SST representation together with the feature’s uncontextualized identity embedding. Both heads are trained with a mean squared error loss on the predicted bin, and the model minimizes their sum over all masked tokens. The feature-level head thereby drives each feature embedding to capture which observed features predict its abundance, while the sample-level head drives the SST to encode the sample’s global abundance structure.

#### Training configuration

Architecture and training hyperparameters were specified a priori, sized for a single consumer GPU with conventional transformer proportions, and used unchanged for every modality. The transformer comprised six layers with a hidden dimension of 256, eight attention heads, a feed-forward dimension of 1,024 and dropout of 0.1. Each sample contributed at most 1,024 detected features, discretized into ten rank-based bins (B 10), with 85% provided as observed context and 15% as reconstruction targets. Both identity-embedding variants were trained per modality, yielding six models. Training used a batch size of 64, a learning rate of 1 × 10⁻⁴, weight decay of 1 × 10⁻⁵ and gradient clipping at 1.0, for up to 200 epochs, retaining the checkpoint with the lowest validation loss. Complete configurations accompany the released checkpoints on Hugging Face.

### 4.3 Attention-derived molecular associations

#### Reference databases

Recovery of molecular relationships was evaluated against nine resources: CORUM (protein complexes)^11^, BioPlex 3.0 (AP–MS co-complex associations)^12^, HuRI (Y2H binary interactions)^13^, STRING (integrated functional associations)^14^, Reactome^15^ and KEGG^16^ (pathway co-membership), GO-CC and GO-BP^17^ (shared cellular-component and biological-process annotation) and OmniPath (transcription factor–target regulation)^18^; all collected 04/26. Each resource was represented as a three-state pairwise label matrix over a shared index of canonical human UniProt accessions: a pair was labelled positive if the relationship was curated, negative if both proteins were represented in the resource but no relationship was annotated between them, and unannotated if either protein was absent. Absence from a resource was thus treated as missing information rather than as evidence against a relationship, whereas annotation was assumed complete for covered proteins. Where required, source identifiers were mapped to canonical UniProt accessions using MyGene.info. Statistics on positive pair counts and filtering steps are reported in Table S3.

#### Enrichment score

This benchmark measures, for each feature, whether its curated partners appear among its highest-scoring candidates more often than chance would place them. The reasoning proceeds in two steps: we first establish how densely true partners populate a random ranking, and then measure how densely they populate the top of the actual ranking. For each feature and reference database, the chance level is the base rate ᵢ *P*ᵢ/ ᵢ, where *P*ᵢ is the number of annotated positive candidates and ᵢ the number of annotated candidates. This is the precision expected from a random ranking. Against it, the candidates of feature were ranked by their pairwise score and the precision at was computed as precᵢ() ᵢ()/ ᵢ(), where ᵢ() is the number of annotated positives and ᵢ() the number of annotated pairs among the top ; unannotated pairs contribute to neither count. Dividing by the base rate gives the enrichment *E*ᵢ() precᵢ()/ ᵢ, so that a random ranking yields one and values above one indicate concentration of true partners near the top. As a single summary, enrichment was averaged over the top 100 candidates per feature and then across all features with at least one annotated pair.

#### Benchmark cohorts

The benchmark was run on the project-held-out test set of each modality (Methods 5.1). For each modality, ten replicates of 1,000 observations were drawn with replacement (seed 21,000 replicate), so that resampling probes robustness to the selected observations. Within each replicate, a common universe was fixed of the 1,000 highest-variance features (after log1p transformation) among those detected in at least 10% of observations, with transcript-level features mapped to their paired canonical UniProt accessions. All methods were scored on this shared universe.

#### Attention read-out

Each evaluated sample was passed ten times through the model with all 1,024 selected input features placed in the observed context, so that every feature attends to every other within a forward pass. Post-softmax attention was averaged across heads within each layer and then across layers, yielding one value per ordered feature pair per pass. Because feature pairs are co-detected at different rates, each pair was averaged only over the passes in which both features were in context. The two directional scores were then averaged into a single symmetric score, with the diagonal left undefined, producing one feature × feature matrix whose row-wise rankings enter the enrichment benchmark.

#### Reference methods

OmicsFM was compared with data-driven association and gene-regulatory inference methods refitted in each replicate (Pearson correlation after total-signal normalization and log1p; FAVA^19^; GENIE3^20^; DeepSEM^21^), a sequence-only control (cosine similarity of ESM-C embeddings) and pretrained single-cell foundation models (scGPT, Geneformer^2^, scPRINT^3^). Hyperparameters are listed in Table S4. The published models take different approaches to network extraction, where one is offered at all: scGPT reads attention from its final layer, which recovered fewer interactions here than layer averaging, and Geneformer exposes no extraction routine. Given this, both were evaluated with the same all-head, all-layer averaging used for OmicsFM. scPRINT was evaluated with its native GNInfer implementation.

#### Gene-universe control

Grounding all references in UniProt protein space could disadvantage the natively gene-level transcriptomic modality and the external single-cell foundation models, either through the feature mapping itself or through reference edges lost in identifier conversion. The single-cell benchmark was therefore repeated entirely in the native gene universe. Attention was extracted from an OmicsFM model trained on the unmapped ENSEMBL gene vocabulary (61,497 genes), the highest-variance feature universe was selected without mapping-based eligibility, and the reference databases were re-indexed to ENSEMBL identifiers and scored directly. The scoring metrics were identical to the main benchmark, but a different resampling design was used: instead of random draws, each of the ten replicates comprised the cells of a single cell type, a regime closer to practical use, where networks are inferred per cell population. Both universes were evaluated on the same replicates. Results are shown in Fig. S2.

#### External BenGRN benchmark

As an externally defined evaluation, scPRINT’s published BenGRN OmniPath benchmark was used with the released v1.6.4 kidney dataset, the o2uniqsx checkpoint and the OmniPath reference bundled with the paper tag. OmicsFM attention was extracted as described above, and both all-pair and transcription-factor-only predictions were scored with the early precision ratio (EPR) and the area under the precision–recall curve (AUPRC), averaged across the ten kidney cell types.

#### Attention vs co-expression

To test whether attention merely recapitulates co-expression, the two were compared directly on the same data. This analysis used the proteomics ESM-C model and its training samples: because the model learned its relational structure from these data, they are the appropriate basis for asking whether that structure reduces to the co-expression they contain. Attention scores were extracted as described above, and co-expression was quantified as the pairwise-complete Pearson correlation between protein log-abundances, using only samples in which both proteins were detected (≥5 co-detections). This co-occurrence requirement matches the model’s own view, which learned to relate two proteins only when they co-occurred in a sample. The attention and co-expression rankings were then compared from both directions: the Pearson correlations of the top 3,000 and 30,000 attention pairs, and conversely the attention scores of the top 3,000 and 30,000 co-expression pairs, were each inspected against the distribution over all pairs.

### 4.4 Tissue-specific attention networks

#### Tissue profiles and network construction

Paired proteome and transcriptome profiles of 30 human tissues were taken from the Wang et al. atlas^23^: proteomics as the authors’ iBAQ quantification, giving a single deep proteome per tissue, and transcriptomics as the paired Human Protein Atlas RNA-seq profiles (v19, E-MTAB-2836) in TPM. Both were mapped onto the UniProt vocabulary. From each tissue profile, attention was read out as described in 5.3 with both the learned-identity and the ESM-C variant of the proteomics and bulk-transcriptomic models, yielding four protein × protein attention matrices per tissue (two modalities × two variants), 116 networks in total.

#### Tissue-specific attention

Testing whether attention reorganizes with tissue identity requires pathways whose activity is restricted to known tissues, together with a control set of processes active in every cell. Both sets were assembled from KEGG using the BRITE hierarchy. The tissue-specific set comprised pathways from the Organismal Systems category, restricted to the six organ-system subcategories (nervous, circulatory, digestive, endocrine, excretory and immune). The housekeeping set, serving as the negative control, comprised pathways from the Genetic Information Processing category (for example, spliceosome, proteasome and DNA replication). Retaining only pathways with at least six members yielded 74 tissue-specific and 24 housekeeping pathways.

Because attention magnitude is not directly comparable across tissues, all protein pairs within a tissue were ranked by attention (rank 1 highest). A pathway’s attention score in a tissue was defined as the geometric mean (GM) of the ranks of its within-pathway pairs, considering only proteins expressed in that tissue so that mere co-occurrence cannot inflate the score. A lower GM thus marks a pathway whose proteins attend to one another more strongly.

For each pathway, the null model of its non-specific behavior was estimated from its GM values across the 30 tissues. These values were approximately normally distributed (Shapiro–Wilk p > 0.05 for 87% of pathways), and because each tested pathway is specific to only a subset of tissues, the bulk of its GM values represents the non-specific background, with tissue-specific values expected only in the left tail. The per-pathway null was therefore modelled from the median and the median absolute deviation (MAD) of its GMs, as both statistics are robust to outliers and hence defined by the non-specific majority. Each observed GM was standardized against this null as z(p,t) (median(GMₚ) − GMₚ,ₜ)/(1.4826 · MADₚ), so that a positive z indicates that pathway p ranks higher in tissue t than in a non-specific tissue. P-values were derived from these z-scores and, because every pathway–tissue cell constitutes a separate test, adjusted with the Benjamini–Hochberg procedure. Tissue-pathway pairs with adjusted p < 0.05 were considered significant. To verify that pathway attention differences were not driven by differences in detectability, the fraction of pathway members expressed in each tissue, corresponding to the proportion of pathway proteins for which an attention score could be computed, was evaluated as a potential confounder for every pathway–tissue pair.

#### Community detection and pathway enrichment

In the preceding analysis, prior knowledge defined the protein clusters and attention was measured within them. This analysis inverts that logic, letting the attention network define clusters without any biological annotation and testing whether these correspond to known pathways. For each tissue, the attention matrix was reduced to a sparse graph retaining the top 3,000 edges, and communities were detected with the Leiden algorithm (resolution 1), discarding communities smaller than twelve proteins. Each community was then tested for enrichment against every KEGG pathway with at least five members in the network, using the hypergeometric test with the network proteins as background, and the pathway with the smallest p-value was assigned as the community label. Because each community is tested against every pathway and only its best match is kept, even a randomly assembled community will show an apparently enriched best match; the observed best-match p-value is therefore optimistically biased. To establish how strong a best match arises by chance alone, an empirical null was estimated: pathway labels were permuted across the network’s proteins 1,000 times, preserving pathway and community sizes, and for every permutation each community was retested against all pathways and its smallest p-value recorded, yielding the distribution of best-match p-values expected for communities with no real pathway coherence. Observed best-match p-values were compared against this null to obtain empirical significance estimates, which were adjusted with the Benjamini–Hochberg procedure across the communities within each tissue.

### 4.5 Sample-level representation evaluation

#### Sample-level benchmark

Evaluating sample-level representations requires more than measuring biological organization within a corpus, because in collections assembled from independent experiments, biological composition and study origin are often confounded, in proteomics especially. A single set of metrics was therefore applied identically to all three modalities, covering three complementary axes: (i) the biological structure captured within each corpus; (ii) the extent to which this structure generalizes across studies; and (iii) the degree to which project identity remains encoded. Biological structure was quantified with the bio-conservation metrics from scIB^24^, excluding cell-cycle, highly-variable-gene and trajectory conservation because they lack comparable counterparts outside single-cell transcriptomics. The remaining six metrics, computed with scIB defaults, were averaged with equal weight into a biological-conservation score (Table 1). Cross-study generalization was evaluated by k-nearest-neighbor label transfer in which all neighbors from the query sample’s own project were excluded, reported as macro-F1. Project-associated structure was assessed by repeating the cluster-based scIB metrics with project identity in place of the biological label; the four resulting metrics (Table 1) were averaged with equal weight into a project-conservation score.

**Table 1.** Metrics used to evaluate sample-level representations. Six scIB-derived metrics quantify biological organization within each corpus and are averaged with equal weight to obtain the biological-conservation score. Cross-project kNN F1 measures biological-label transfer using only neighbors from other projects. Four corresponding metrics quantify project-associated structure and are averaged with equal weight to obtain the project-conservation score.

| Evaluation axis | Metric | Interpretation |
| --- | --- | --- |
| Biological organization | Label silhouette | Compactness and separation of samples sharing the same biological label |
|  | Leiden NMI | Information about biological labels preserved by unsupervised Leiden clusters |
|  | Leiden ARI | Chance-corrected agreement between Leiden clusters and biological labels |
|  | Isolated-label silhouette | Separation of biological labels represented in relatively few projects |
|  | Isolated-label F1 | Recovery of labels represented in relatively few projects as distinct Leiden clusters |
|  | Graph cLISI | Preservation of biological-label structure in local neighborhoods |
| Cross-project generalization | Cross-project kNN F1 | Macro-F1 of biological-label transfer using only neighbors from other projects |
| Project-associated structure | Project silhouette | Compactness and separation of samples originating from the same project |
|  | Leiden NMI, project | Information about project identity preserved by unsupervised Leiden clusters |
|  | Leiden ARI, project | Chance-corrected agreement between Leiden clusters and project identity |
|  | 1 – Graph iLISI | Local homogeneity of project identities in the neighborhood graph |

Evaluation cohorts were constructed identically for the three modalities. For the main benchmark, the training, validation and test partitions were pooled within each modality. This was particularly important for proteomics, whose test partition contains only 769 eligible samples across nine tissues, substantially limiting biological coverage. From each pooled corpus, ten deterministic cohorts of 5,000 observations were drawn (seeds 21,000–21,009). Cohort allocation was first distributed as evenly as possible across eligible biological labels, subject to availability. Within each label, observations were selected round-robin across the contributing projects, drawing without replacement from each label–project group, so that highly prevalent labels and large projects could not dominate. All representation methods were evaluated on identical observations within each cohort. Performance is reported per modality as the mean and standard deviation across the ten cohorts.

OmicsFM SST embeddings from both variants were compared with PCA, an autoencoder and log-transformed raw abundances, and, for single-cell data, additionally with scGPT, Geneformer and scPRINT. PCA and the autoencoder were fitted once per modality on the cohort union, ensuring that between-cohort variation reflects cohort composition rather than model refitting, and all pretrained models were applied without fine-tuning. A random embedding, drawn independently of the data, provided the lower bound that each metric returns in the absence of any structure. Because the main benchmark pools all partitions, OmicsFM was evaluated in part on observations seen during pretraining, whereas PCA and the autoencoder were fitted on the evaluation cohorts themselves; the evaluation was therefore repeated on held-out test data alone (Fig. S7), using all 769 eligible proteomics test samples and, for bulk and single-cell transcriptomics, one deterministic label- and project-balanced cohort of 5,000 observations drawn from each test set with the same hierarchical procedure.

#### Robustness to protein dropout

Public proteomic profiles vary widely in coverage, and because the SST is pretrained on randomly subsampled protein sets, its representations may inherit robustness to missing features. We tested this directly by perturbing held-out profiles. The analysis used the proteomics model and its test set. For each sample, detected proteins were randomly removed at dropout levels from 0% to 100%, with five independent masks per non-zero level. Perturbed profiles were then represented in three ways: with the SST, with a dimension-matched 256-component PCA and with the full-dimensional log-transformed abundance vector. PCA was fitted once, on the unperturbed profiles, and because each input-collator draw samples up to 1,024 detected proteins, SST representations were averaged over five draws.

Representation stability was quantified as excess self-correlation, which asks how similar a sample’s perturbed representation remains to its own unperturbed one, beyond the similarity any two samples share in that space. For each sample and dropout level, the Pearson correlation between the perturbed and unperturbed representations was computed after centering the representation space on the mean of the unperturbed cohort. From this self-correlation, the sample’s mean correlation with the complete reference cohort was subtracted. These corrections remove the correlation that the overall geometry of a representation space produces on its own, so that the excess reflects retained sample identity alone.

Preservation of biological organization was assessed by cross-project tissue retrieval, which asks whether a perturbed sample still lies closest to samples of its own tissue from other studies. Perturbed samples were queried against the fixed unperturbed reference set using Euclidean distance, excluding references from the query’s own project, and the AUROC measured whether same-tissue references ranked closer than different-tissue ones. To express performance on a scale where 0 denotes chance and 1 perfect recovery, the AUROC was normalized against a shuffled baseline, obtained at each dropout level by permuting the labels of the perturbed samples while leaving the reference set unchanged: AUROC_normalized (AUROC_observed − AUROC_shuffled)/(1 − AUROC_shuffled). This evaluation included all samples from tissues represented in at least two projects, 622 samples across seven tissues. Results are reported as means across each dropout level, with 95% percentile bootstrap confidence intervals from 1,000 sample-level resamples.

### 4.6 Downstream applications

#### Cell-type classification from single-cell proteomics

Transfer of the sample-level representation was tested on cell-type classification. PXD071075 contains single-cell proteomic profiles from three developing human brain donors sampled at gestational weeks 13, 15 and 19^25^. Because each donor represents one developmental stage, donor and stage effects are confounded, and evaluation on the held-out donor therefore probes a combined donor and developmental-stage shift. PXD071075 was for the evaluated OmicsFM model explicitly removed from the pretraining corpus. An MLP classification head was attached to the SST, and three configurations were compared: end-to-end fine-tuning, an MLP trained on frozen pretrained representations, and an MLP trained on representations from a frozen encoder initialized without pretraining. L1- and L2-regularized logistic regression and a random forest, trained on log₂(x 1)-transformed protein abundances, served as conventional baselines. Hyperparameters for the OmicsFM configurations were selected by training on GW15 and ranking performance on GW13 by mean macro-F1 across three repeats. The selected configurations were then retrained on GW13 and GW15 and evaluated on held-out GW19 in three fresh repeats. Performance was measured as accuracy and macro-F1, with 95% percentile confidence intervals from 1,000 bootstrap resamples of the GW19 cells.

#### Gene-essentiality prediction

This application probes the learned feature-identity embeddings of OmicsFM, asking how much functional information they encode about each gene. General gene essentiality was derived from the DepMap Public 24Q4 CRISPR GeneEffect dataset^26^: for each gene, all finite Chronos scores^27^ were averaged across 1,178 cancer cell lines, yielding one essentiality score for each of 17,692 genes. More-negative scores indicate greater loss of fitness upon knockout. Inputs comprised frozen ESM-C protein-sequence embeddings and the learned feature-identity embeddings of the (learned-identity) OmicsFM models from proteomics, bulk transcriptomics and single-cell transcriptomics, evaluated individually and as pairwise or complete concatenations. Two matched controls tested whether gains from concatenation could arise from increased input dimensionality alone: ESM-C concatenated with Gaussian noise features of matching dimension (C1), and ESM-C concatenated with OmicsFM embeddings randomly permuted across genes (C2), which preserves the embedding distribution while destroying gene correspondence.

For each input representation, a multilayer perceptron was trained under nested five-fold gene-level cross-validation. In each outer fold, 20% of genes were held out for testing, and the remaining genes were divided into inner training and validation sets for selecting the network architecture, dropout, learning rate and stopping epoch. Models were optimized with AdamW and a Huber loss, refitted on the complete outer-training fold, and predictions from three random seeds were averaged. Performance was evaluated on the combined out-of-fold predictions with Pearson correlation as the primary metric.

#### In-silico perturbation prediction

We evaluated perturbation-response prediction on three CRISPR screens: Replogle K562 (1,822 single-target perturbations)^28^, Adamson (86 perturbations)^29^, and Norman (283 conditions, including 131 two-gene combinations)^30^. Each dataset was partitioned into five cross-validation folds at the perturbation-condition level. All cells belonging to the same perturbation were kept together, and every perturbation appeared in the test set exactly once. Within each fold, the remaining conditions were divided into training (90%) and validation (10%) sets. For Norman, splits were constructed using order-independent target sets, ensuring that alternative label orientations of the same gene combination remained in the same partition. All methods were trained and evaluated on a shared protein-coding gene universe comprising 5,757, 3,775, and 3,212 genes for Replogle, Adamson, and Norman, respectively.

Perturbation responses in these screens contain a substantial generic component shared across conditions. Consequently, even a baseline that predicts the same response for every perturbation can achieve a high correlation^37^. To isolate condition-specific effects, we first defined the total response of each perturbation as the difference between its mean expression profile and that of the control cells. We then estimated a generic response vector as the mean total response across all training perturbations. Averaging across many perturbations is expected to attenuate condition-specific effects while retaining the response shared across conditions.

The specific response of a condition was therefore defined as its total response minus this generic response, which was estimated exclusively from the training data.

For each held-out perturbation, we calculated the Pearson correlation between the predicted and observed specific-response vectors. Correlations were computed across either all evaluated genes, termed specific Pearson delta, or the 20 genes with the largest absolute observed specific responses for that condition, termed specific Pearson delta DE20. Because Pearson correlation is insensitive to scale, we additionally report the amplitude ratio between the predicted and observed responses to assess whether the predicted effects have the correct magnitude.

We estimated the reliability ceiling of these correlation metrics from the internal agreement of the experimental data. For each held-out perturbation, its cells were randomly divided into two equally sized groups, and the mean specific-response vector was calculated for each half. The two vectors were then correlated across genes, and the resulting split-half correlations were averaged across perturbations. Because the benchmark response was calculated using all available cells rather than half of the cells, we converted the mean split-half correlation to full-sample reliability using the Spearman-Brown correction. A perfect model predicts the underlying biological response but is evaluated against a noisy experimental mean rather than the true response. We therefore defined the maximum expected model correlation as the square root of the corrected full-sample reliability: ceiling = 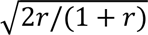, where *r* is the mean split-half correlation across perturbations.

OmicsFM (ESM-C) was fine-tuned independently from each of its three pretrained checkpoints, corresponding to proteomics, bulk transcriptomics, and single-cell transcriptomics. Fine-tuning used a delta-MSE objective between the predicted and observed expression changes relative to control. As a pretraining ablation, the same architecture was also trained from random initialization. scGPT was initialized from the whole-human checkpoint and fine-tuned separately within each fold following its official perturbation-prediction tutorial. GEARS was trained using the official implementation and default hyperparameters. The linear baseline was adapted from Ahlmann-Eltze et al.^37^. It models perturbation responses using bilinear ridge regression over 10-dimensional PCA gene embeddings and represents an unseen perturbation using the embedding of its target gene.

## Supporting information

Supplementary Data 1. ProteomeXchange accessions of the 1,143 public proteomics datasets reprocessed for the OmicsFM pretraining corpus

## Data and code availability

The OmicsFM source code is available at https://github.com/CompOmics/OmicsFM. Pretrained model checkpoints for all three modalities are available on Hugging Face at https://huggingface.co/rednaSander/omicsfm. The bulk and single-cell transcriptomic training corpora are available at https://huggingface.co/datasets/rednaSander/omicsfm-data; the reprocessed proteomics corpus will be made available in the same repository upon publication. The tissue-specific attention networks (30 human tissues, proteomics and transcriptomics) were deposited to Zenodo (https://doi.org/10.5281/zenodo.22069867). The dataset-construction pipelines (the run-level metadata-annotation pipeline with its resulting annotations, and the transcriptomics corpus builders) were also deposited to Zenodo (https://doi.org/10.5281/zenodo.22071865), as well as the code, inputs and outputs of all manuscript experiments at https://doi.org/10.5281/zenodo.22072026. All public datasets used are referenced in supplementary data 1, under their original accessions.

## Acknowledgements

T.C. and R.G. acknowledge funding from the Research Foundation Flanders (FWO) [12A8W25N, 12AK526N]. L.M. acknowledges funding from the Horizon Europe Project COMBINE [101191739 and the CHIST-ERA project ODEEP-EU [G0GDV23N]. We thank all laboratories that deposited their data in ProteomeXchange, GEO, and the CELLxGENE Census; this work would not have been possible without public data sharing. The complete list of the 1,143 reprocessed ProteomeXchange projects that constitute the proteomics corpus is provided in Supplementary Data 1.

## Supplementary information

**Figure S1.**
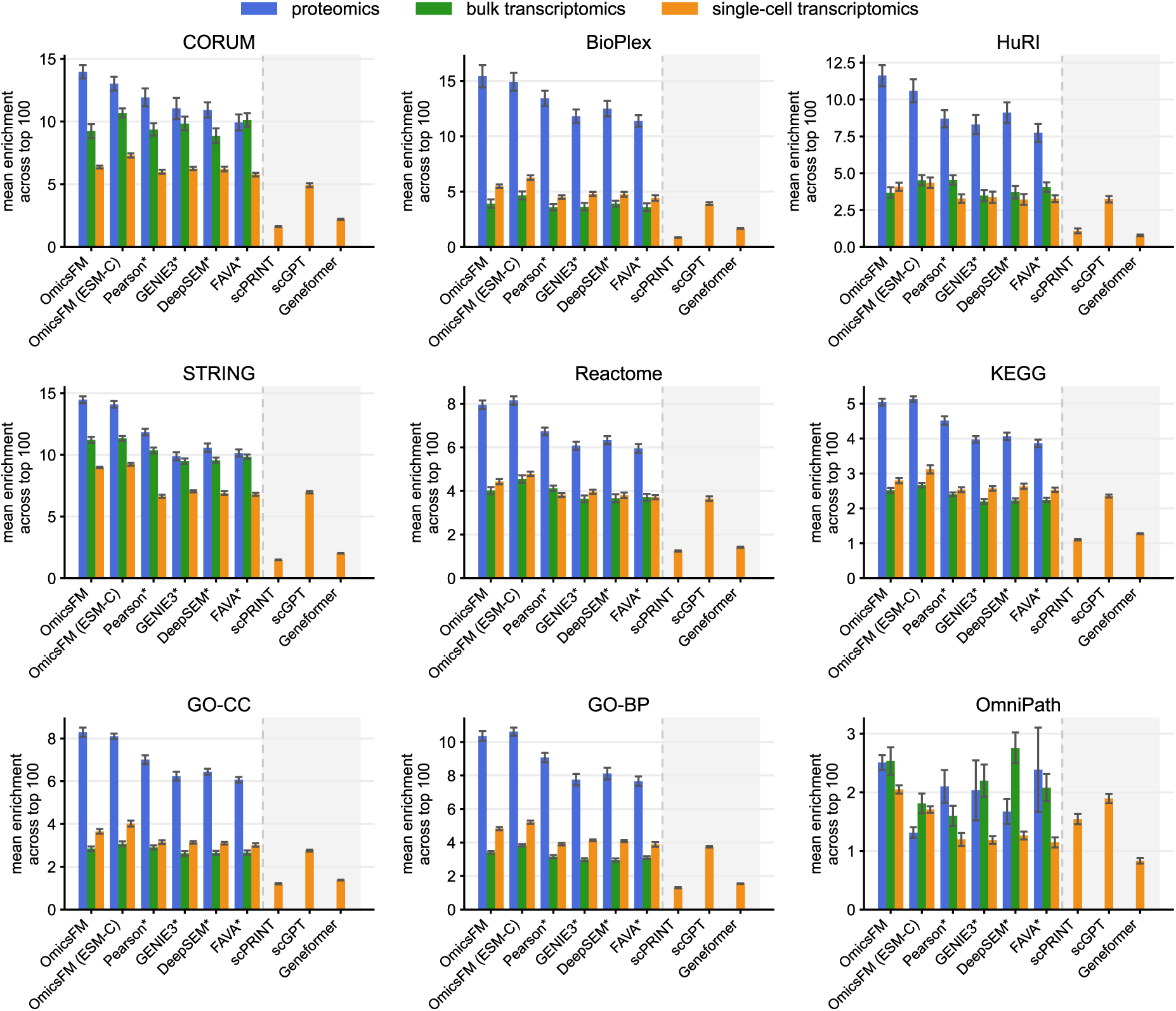
Interaction recovery across nine evidence databases and three modalities. Each panel is one evidence database, bars give the mean enrichment across the top 100 ranked pairs (random 1) for proteomics (blue), bulk transcriptomics (green) and single-cell transcriptomics (orange). Every method is evaluated on the same 10 random 1,000-observation draws from the project-held-out test split, with the top-1,000 HVGs reselected inside each draw. Bar height is the mean enrichment over those 10 draws and the error bar is their standard deviation. That spread is small relative to the between-method differences for every database except OmniPath, where the error bars overlap across most methods. Methods marked with an asterisk (*) are fit directly on each replicate, whereas OmicsFM is applied zero-shot, with no exposure to the evaluation draw. The starred methods therefore carry a systematic advantage, and part of their variance reflects refitting on each draw as well as the draw itself.

**Figure S2.**
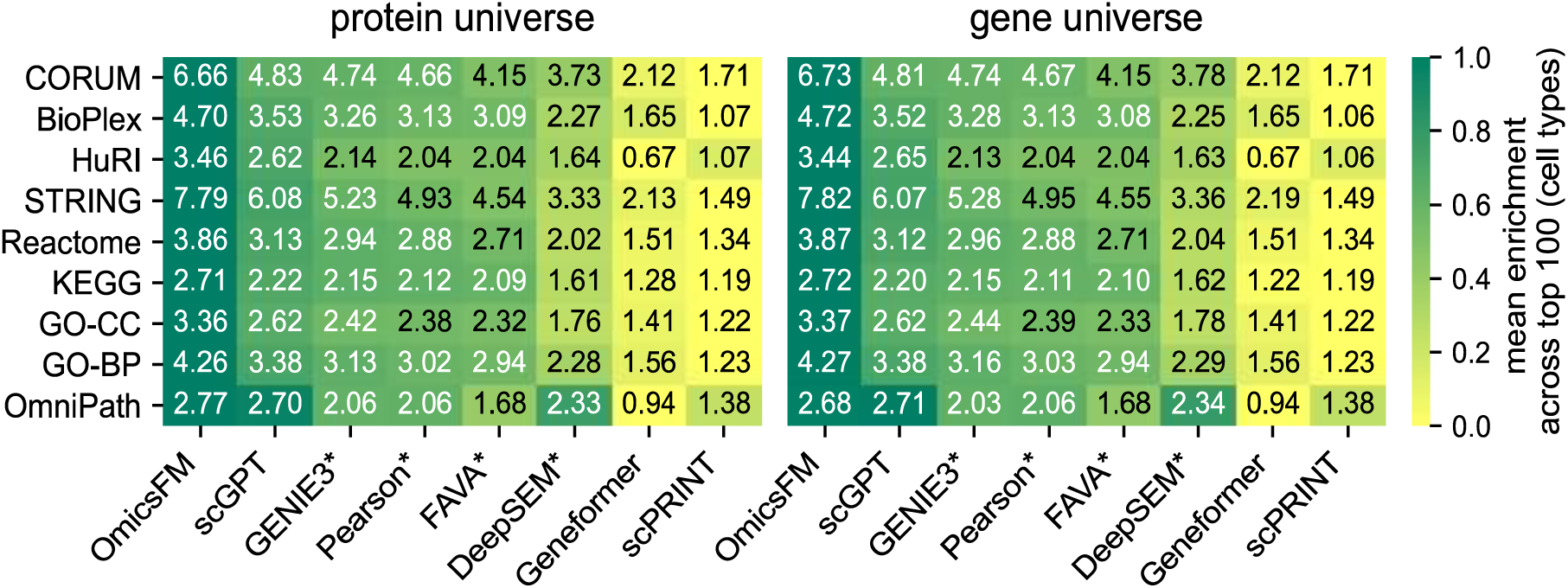
Grounding the association benchmark in UniProt protein space does not disadvantage gene-level methods. The single-cell benchmark was run in the protein universe (left) and repeated entirely in the native gene universe (right), in which OmicsFM attention derives from a model trained on the unmapped ENSEMBL vocabulary and all references are re-indexed to genes without a mapping step (Methods 5.3). Each of the ten replicates comprised the cells of a single cell type, and both universes were evaluated on identical replicates, so the panels differ only in feature universe. Printed values give the mean enrichment over the top 100 ranked partners per feature, averaged across the ten cell types; methods marked with an asterisk* were refitted per replicate. Enrichment differs by at most 0.09 between universes for every method and database, and the method ranking is unchanged. Reference edges lost in identifier conversion are quantified in Table S3.

**Figure S3.**
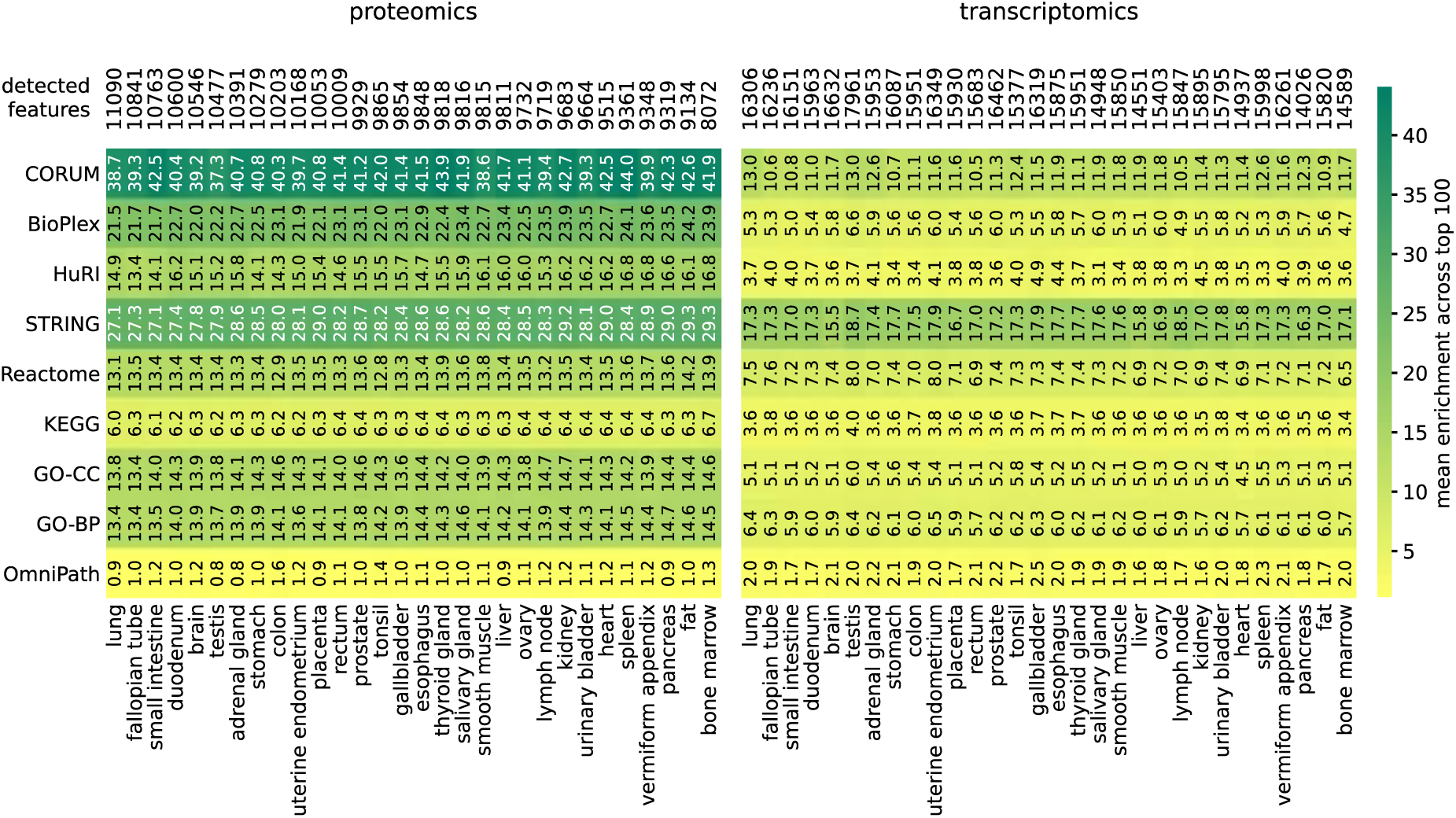
Enrichment of curated molecular relationships in the tissue-specific attention networks. Attention networks were derived from the deep proteome and transcriptome profiles of 30 human tissues [cite] with the benchmark-best variant per modality (learned-identity variant for both proteomics and bulk transcriptomics; Methods 5.4). For each tissue (columns) and reference resource (rows), the printed value and color give the mean enrichment over the top 100 ranked partners per protein: at each rank k, the precision among a protein’s k highest-attention annotated partners is divided by that protein’s base rate of true partners in the resource, and enrichment is averaged over ranks 1–100 per protein and then over all proteins with at least one annotated pair (Methods 5.3). A value of 1 corresponds to a random ranking. The top row lists the number of detected features in each tissue profile. Unlike the held-out benchmark (Fig. 2A), which is restricted to a shared 1,000-feature universe, these networks span all detected proteins; at this broader scale, the proteomics advantage for complex-, interaction- and pathway-based resources becomes more pronounced, whereas OmniPath transcription-factor–target enrichment remains higher in transcriptomics.

**Figure S4.**
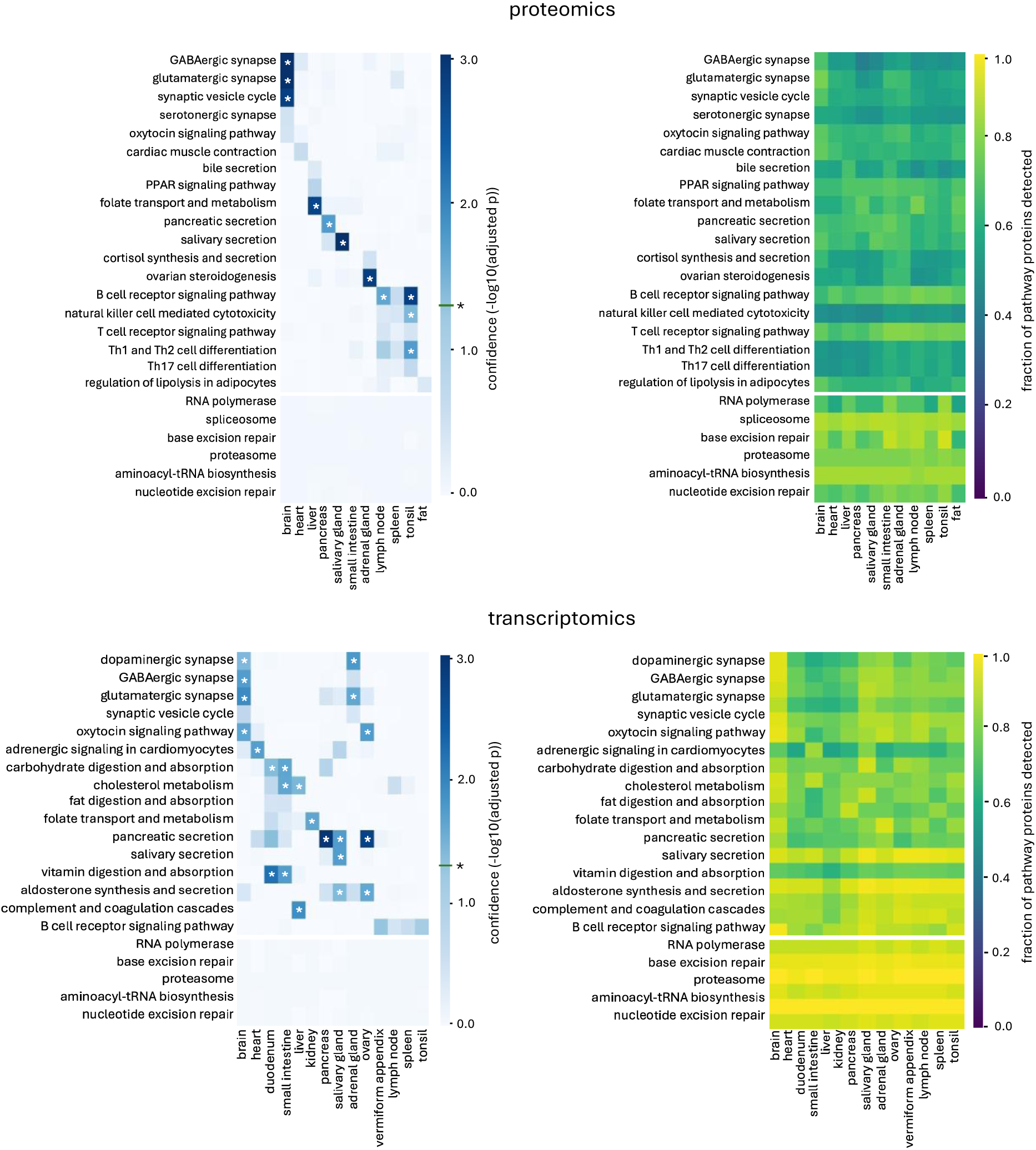
Tissue-specificity of pathway attention and its detectability control from proteomics and transcriptomics tissue-specific attention networks. Within-pathway attention was scored per pathway and tissue as the geometric mean rank of within-pathway pairs and standardized against a robust per-pathway null estimated across the 30 tissues (Methods 5.4). Top row: proteomics (learned-identity variant); bottom row: bulk transcriptomics (learned-identity variant). Left panels: confidence of tissue-specific pathway attention as the −log10(adjusted p); the level marked with an asterisk* on the color bar indicates the significance threshold (adjusted p 0.05), and asterisks in cells mark significant pathway–tissue pairs. Rows are split into tissue-specialized pathways (upper block) and housekeeping pathways (lower block), which serve as the negative control; columns show tissues. Right panels: the corresponding detectability control, giving for each pathway–tissue pair the fraction of pathway proteins detected in that tissue’s profile, and hence the proportion for which attention scores could be computed. Tissue-restricted pathways reach significance in their expected tissues, whereas housekeeping pathways show no tissue preference; because the detection fractions are comparatively uniform across tissues, the significant attention differences are not explained by pathway detectability.

**Figure S5.**
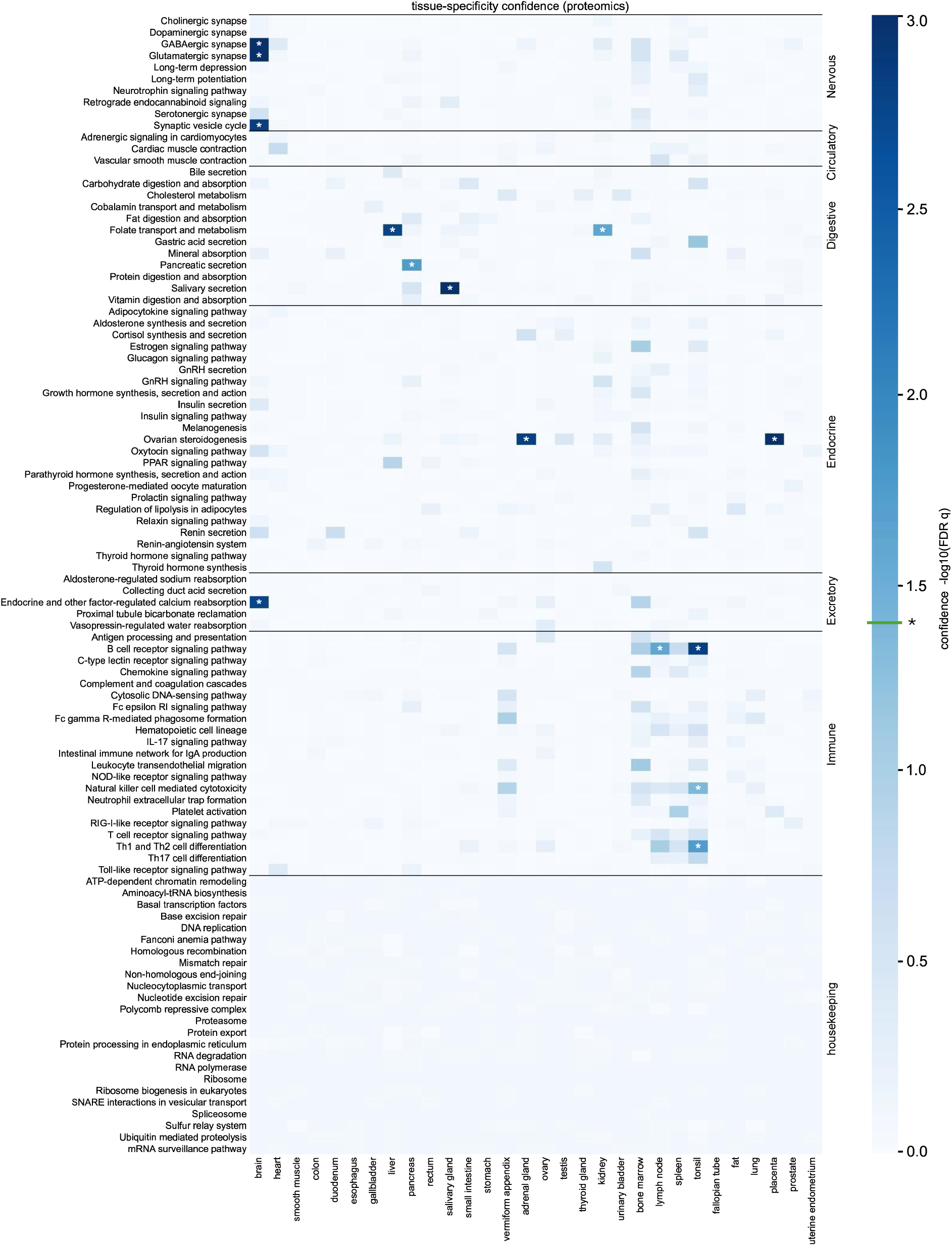
Tissue-specificity of KEGG pathway attention scores (Proteomics). Full Heatmap of confidence (−log10 FDR-adjusted p) that a pathway is more strongly activated in a given tissue than in the typical tissue, across all 30 tissues (columns, grouped by organ system) and all tested KEGG pathways (rows). Rows are ordered by organismal-system annotation (nervous, circulatory, digestive, endocrine, excretory, immune), followed by housekeeping negative controls; black lines separate the blocks. Per pathway, tissue scores were converted to robust z-scores (median/MAD) and tested against a standard normal, with Benjamini–Hochberg correction applied separately within the tissue-specific and housekeeping families; Asterisks* mark cells with FDR < 0.05.

**Figure S6.**
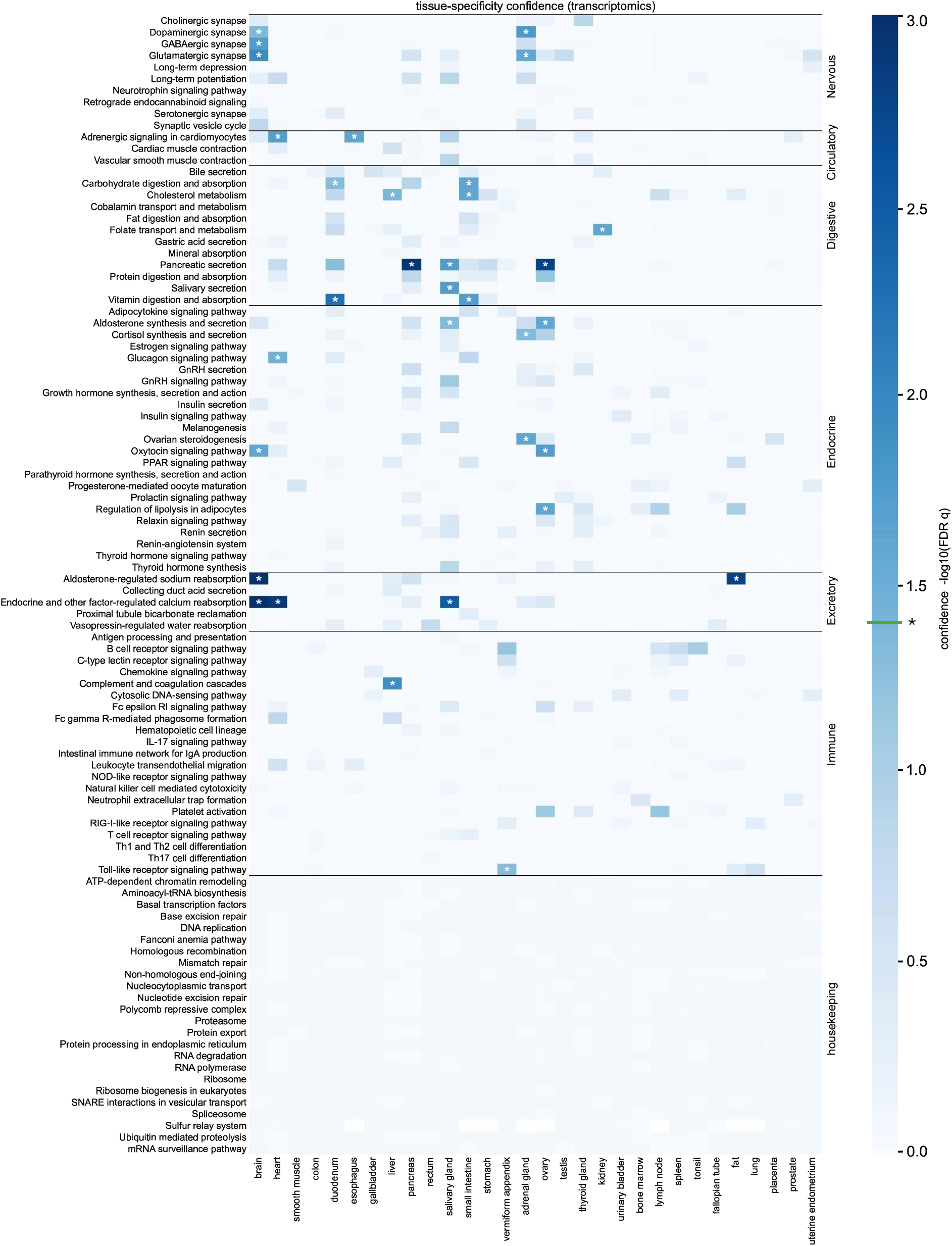
Tissue-specificity of KEGG pathway attention scores (bulk transcriptomics). Full Heatmap of confidence (−log10 FDR-adjusted p) that a pathway is more strongly activated in a given tissue than in the typical tissue, across all 30 tissues (columns, grouped by organ system) and all tested KEGG pathways (rows). Rows are ordered by organismal-system annotation (nervous, circulatory, digestive, endocrine, excretory, immune), followed by housekeeping negative controls; black lines separate the blocks. Per pathway, tissue scores were converted to robust z-scores (median/MAD) and tested against a standard normal, with Benjamini–Hochberg correction applied separately within the tissue-specific and housekeeping families; Asterisks* mark cells with FDR < 0.05.

**Figure S7.**
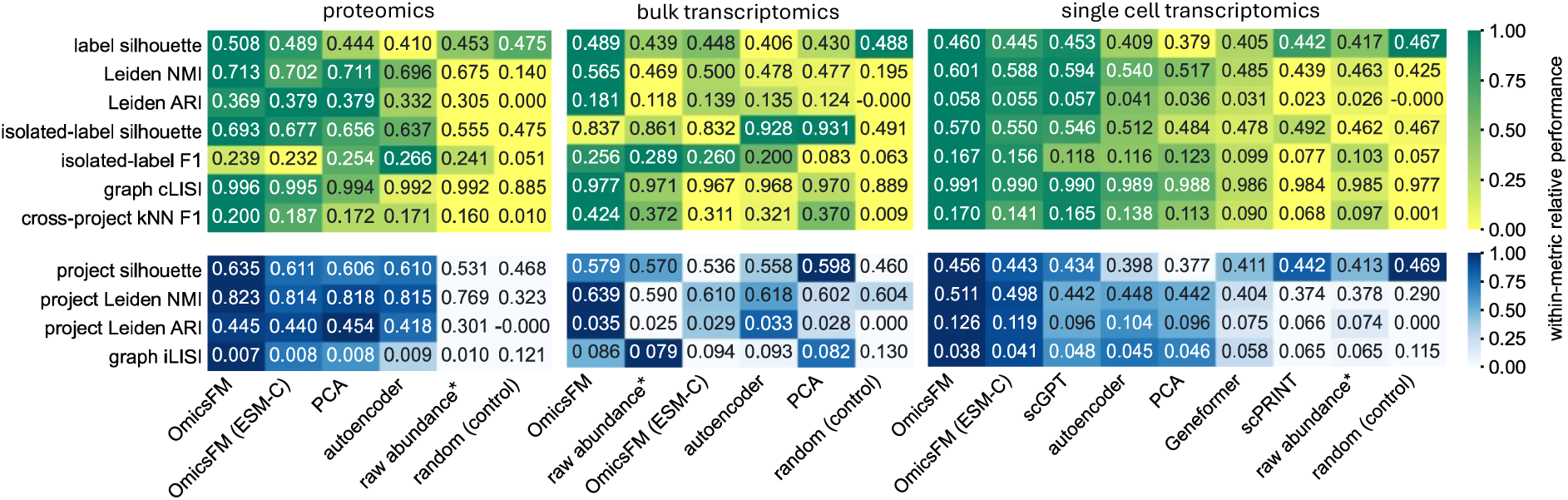
Individual metric scores underlying the aggregated sample-level benchmark. For each modality, rows show the individual metrics entering the biological-conservation score (upper block, shown together with cross-project kNN F1) and the project-conservation score (lower block); columns show the compared representations ordered from average highest to lowest score across all metrics. Printed values are mean scores across the ten label- and project-balanced evaluation cohorts (Methods 5.5). Cell color encodes performance relative to the other representations within each metric and modality, so colors are comparable within a row but not across rows or modalities. Raw abundance, marked with an asterisk, is not a learned representation and retains the full protein space; the random embedding provides the lower bound each metric returns in the absence of structure. Cohort-level variability for these scores is shown in Fig. S6.

**Figure S8.**
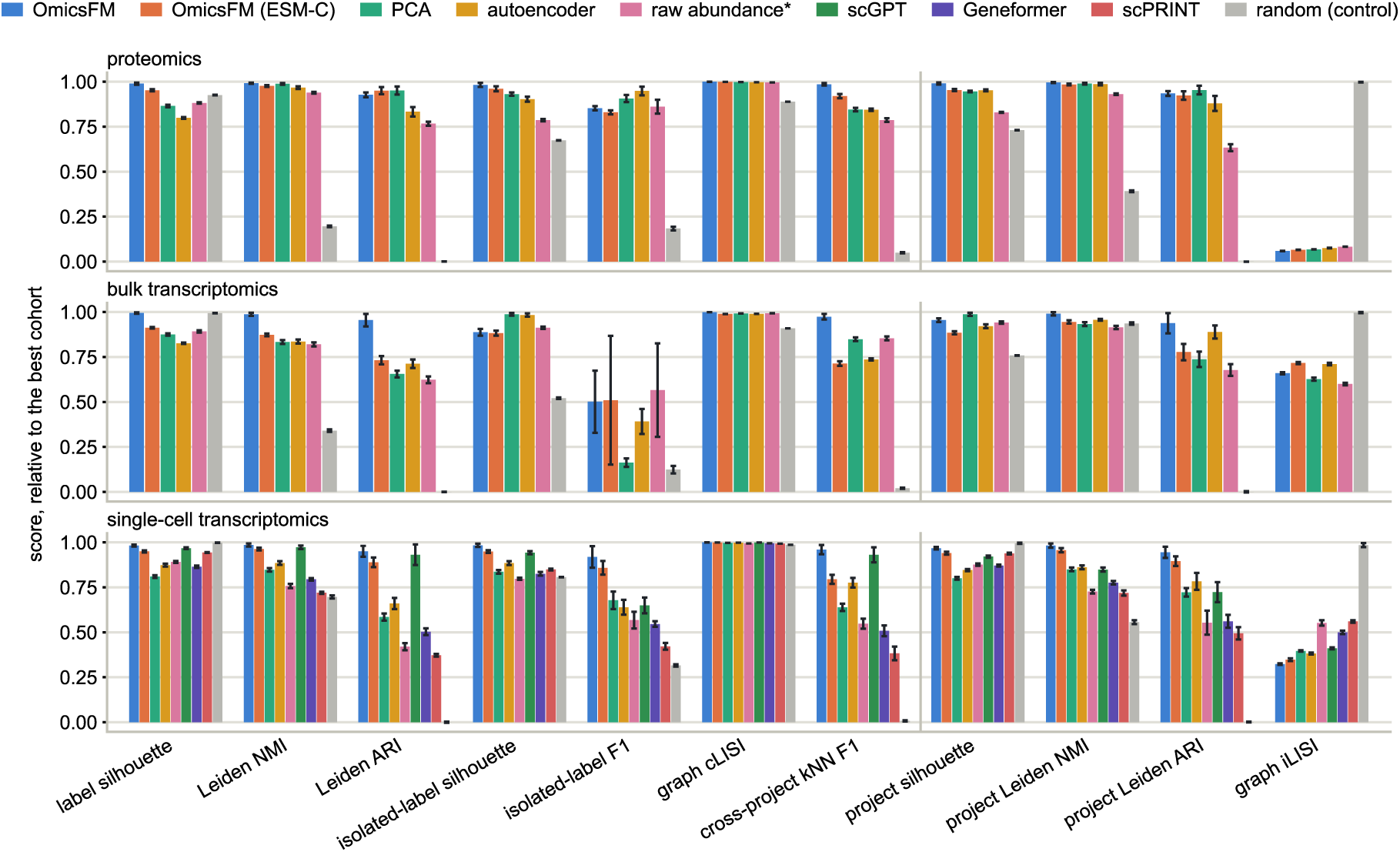
Cohort-level variability of the individual metric scores. The metrics and representations of Fig. S5, shown per modality with their variation across the ten evaluation cohorts. For each metric, scores were normalized to the best score achieved by any representation in any cohort, so that 1 denotes the overall best performance observed for that metric. Bars show the mean of this relative score across the ten cohorts, and error bars its standard deviation. Bio-conservation metrics and cross-project kNN F1 are grouped left of the divider and project-conservation metrics right of it. The legend within each panel lists representations in order of overall performance. Raw abundance, marked with an asterisk*, is not a learned representation. The random control provides the no-structure reference.

**Figure S9.**
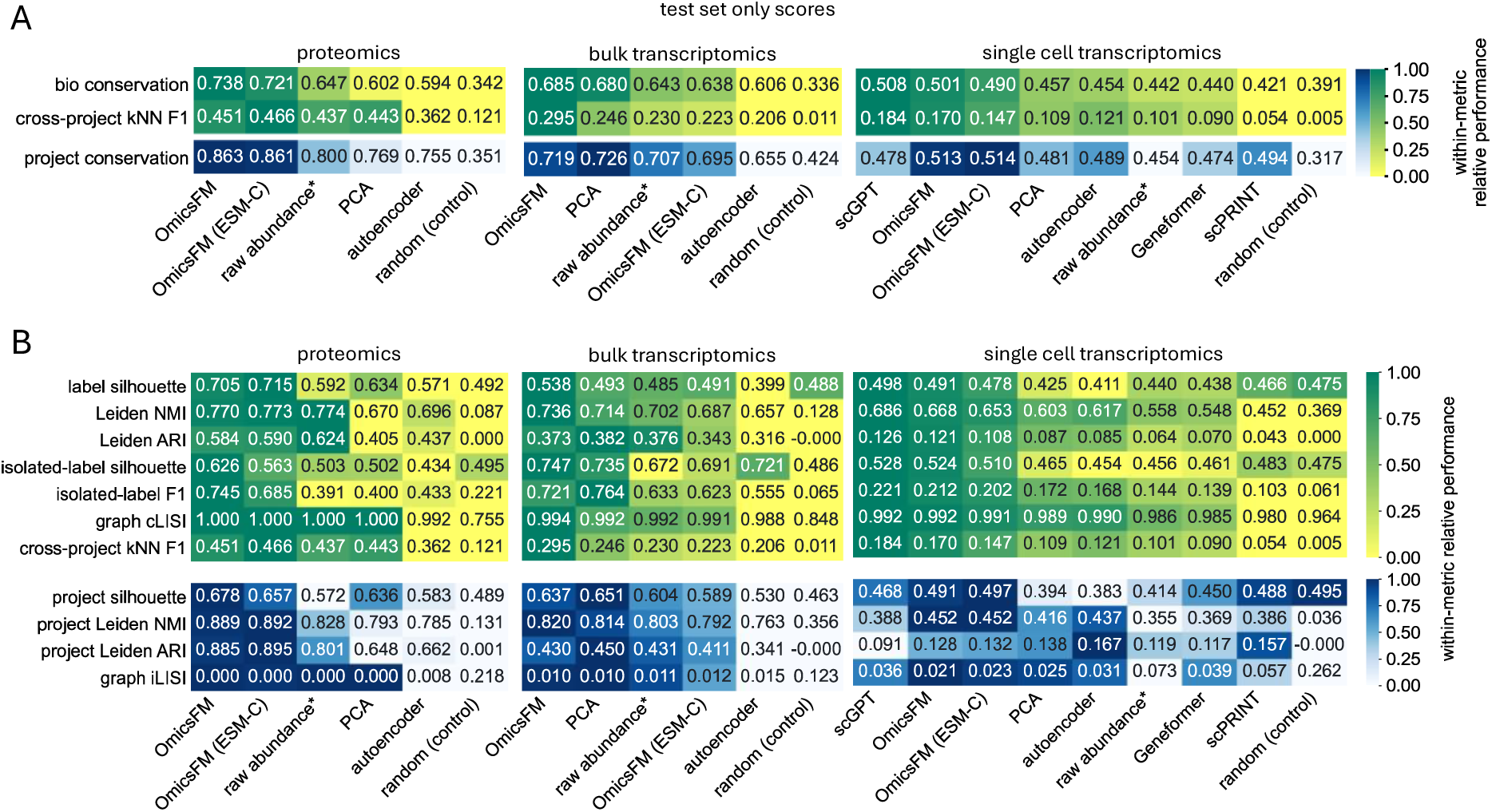
Sample-level benchmark restricted to the project-held-out test sets. Because the main benchmark pools all data partitions, the evaluation was repeated on held-out test data alone (Methods 5.5), using all 769 eligible proteomics test samples and one deterministic label- and project-balanced cohort of 5,000 observations drawn from each transcriptomic test set. Every observation was unseen during OmicsFM pretraining; no equivalent guarantee holds for the external single-cell foundation models, whose pretraining corpora may overlap with the evaluated datasets. **(A)** Aggregated scores per modality: bio conservation (mean of the six scIB metrics), cross-project kNN F1 and project conservation (mean of the four project metrics). **(B)** The individual metrics underlying the aggregates, with the bio-conservation metrics and cross-project kNN F1 in the upper block and the project-conservation metrics in the lower block. Printed values are the scores on the single test cohort per modality; because only one cohort is evaluated, no across-cohort variability is available, in contrast to Figs. S5 and S6. Cell color encodes performance relative to the other representations within each metric and modality, so colors are comparable within a row but not across rows. Columns are ordered by highest average score within each modality. Raw abundance, marked with an asterisk*, is not a learned representation and retains the full feature space; the random embedding provides the no-structure lower bound.

**Table S1.** Coverage of the run-level metadata annotations in the proteomics corpus. For each annotated entity, the table lists the number and percentage of the 48,837 corpus samples with a non-empty value after source merging and ontology normalization, together with the number of unique values. Because a sample originates from either a tissue or a cell line, these two fields are mutually exclusive: the second percentage for each therefore gives coverage among the samples for which the field is applicable, excluding those annotated with the other field. The first row reports project and run identifiers, covering 1,143 ProteomeXchange projects and 48,012 unique run names.

| entity | number of values | % of annotation | unique values |
| --- | --- | --- | --- |
| pxd, run | 48,837 | 100.0 | 1,143 / 48,012 |
| instrument | 43,343 | 88.8 | 70 |
| acquisition | 42,157 | 86.3 | 3 |
| organism | 41,647 | 85.3 | 16 |
| modifications | 40,592 | 83.1 | 31 |
| labeling | 40,382 | 82.7 | 7 |
| enzymes | 40,086 | 82.1 | 12 |
| lc_column | 36,823 | 75.4 | 6 |
| fragmentation | 30,693 | 62.8 | 11 |
| ionization | 30,051 | 61.5 | 4 |
| tissue | 24,314 | 49.8 → 63.6 | 138 |
| disease | 24,262 | 49.7 | 124 |
| fractionation | 23,473 | 48.1 | 287 |
| treatment_class | 16,079 | 32.9 | 7 |
| cell_line | 15,328 | 31.4 → 50.3 | 102 |
| enrichment | 11,656 | 23.9 | 211 |
| cell_part | 7,214 | 14.8 | 24 |
| collision_energy | 183 | 0.4 | 2 |

**Table S2.** Field-level performance of the manuscript-extraction pipeline on the 30 held-out manuscripts. For each extracted metadata field, grouped by category, precision, recall and F1 were computed against the manually curated ground-truth annotations. Manuscripts (n) gives the number of held-out manuscripts in which the field was annotated and therefore contributed to the scores.

| Category | extracted field | precision | recall | F1 | manuscripts, n |
| --- | --- | --- | --- | --- | --- |
| sample biology | organism | 0.983 | 1.000 | 0.989 | 30 |
|  | tissue | 0.805 | 0.770 | 0.782 | 29 |
|  | disease | 0.774 | 0.747 | 0.756 | 28 |
|  | cell part | 0.500 | 0.500 | 0.500 | 2 |
|  | cell line | 0.667 | 0.667 | 0.667 | 9 |
| sample preparation | enzymes | 0.967 | 0.933 | 0.944 | 30 |
|  | modifications | 0.947 | 0.803 | 0.813 | 30 |
|  | labelling | 0.983 | 1.000 | 0.989 | 30 |
|  | fractionation | 0.767 | 0.756 | 0.760 | 30 |
|  | enrichment | 0.417 | 0.417 | 0.417 | 6 |
| MS configuration | instrument | 0.983 | 0.978 | 0.972 | 30 |
|  | fragmentation | 1.000 | 1.000 | 1.000 | 29 |
|  | collision energy | 0.788 | 0.808 | 0.795 | 26 |
|  | acquisition mode | 1.000 | 0.983 | 0.989 | 30 |
|  | LC column | 0.967 | 0.967 | 0.967 | 30 |
|  | gradient time | 0.879 | 0.879 | 0.874 | 29 |
|  | ionization | 0.357 | 0.357 | 0.357 | 28 |

**Table S3.** Retention of curated positive pair labels after resource-specific filtering and mapping to canonical human UniProt accessions. For each reference resource, the table lists the filtering criteria applied before identifier conversion, followed by the number of positive pairs before mapping. It then reports pairs removed because one or more native identifiers lacked a UniProt counterpart, pairs that became redundant when multiple native identifiers mapped to the same protein, the final retained positive-pair count, and the percentage lost. Mapping preserved all positive pairs for seven resources; only HuRI (1.71%) and KEGG (8.14%) showed appreciable loss.

| database | filtering | unmappable | mapping many to one | edges after mapping | % lost by mapping |
| --- | --- | --- | --- | --- | --- |
| CORUM | restricted to $\geq 2$ subunits | 0 | 0 | 47,943 | 0.00% |
| BioPlex | probability $\geq 0.75$ & isoform collapsed to canonicals | 0 | 0 | 108,968 | 0.00% |
| HuRi | all included | 890 | 240 | 50,938 | 1.71% |
| STRING | restricted confidence $\geq 700$ | 0 | 0 | 224,574 | 0.00% |
| Reactome | restricted to human | 0 | 0 | 999,227 | 0.00% |
| KEGG | restricted to human | 264,364 | 64,645 | 2,917,497 | 8.14% |
| GO-CC | only experimental & manually | 0 | 0 | 1,685,546 | 0.00% |
| GO-BP | only experimental & manually | 0 | 0 | 812,068 | 0.00% |
| OmniPath | restricted to TF-gene pairs | 0 | 0 | 113,686 | 0.00% |

**Table S4.** Hyperparameter settings for the reference network-inference methods. (GENIE3, DeepSEM, and FAVA) used in the attention-based network recovery benchmark. For all tools the authors’ default parameters were used.

| method | hyperparameter | value used | source / note |
| --- | --- | --- | --- |
| GENIE3 | tree ensemble | Extra-Trees (ExtraTreesRegressor) | chosen for stabler importances on correlated regulators |
|  | number of trees | 1000 | author default |
| | max_features | $\sqrt{p}$ ($\approx 32$ of 1,000 HVG) | author default |
| DeepSEM | epochs | 90 cell type non-specific;<br>150 cell type specific | author default |
|  | batch size | 64 | author default |
| | learning rate | $1 \times 10^{-4}$ (RMSprop) | author default |
| | $\alpha$ (L1 weight on W) | 100 cell type non-specific;<br>1 cell type specific | author default |
| | $\beta$ (KL weight) | 1 cell type non-specific;<br>0.01 cell type specific | author default |
|  | hidden units | 128 | author default |
|  | alternating opt. (K1, K2) | 1, 2 | author default |
| | LR decay ($\gamma$) | 0.99 | author default |
| FAVA | hidden layer | 1,000 units | author default |
|  | latent dimension | 100 | author default |
|  | epochs | 50 | author default |
|  | batch size | 32 | author default |
| | learning rate | $1 \times 10^{-3}$ (Adam, clipnorm $1 \times 10^{-3}$) | author default |
|  | loss weighting | 0.9-recon + 0.1-KL | author default |

